# *Enterococcus faecalis* biofilm rewires neutrophil metabolism to suppress antimicrobial activity

**DOI:** 10.64898/2026.06.18.733107

**Authors:** Haris Antypas, Ronni Anderson Gonçalves Da Silva, Geneve Kang Xin Wong, Cui Liang, Logeshwari Muthualagu Natarajan, Rachel Jing Wen Tan, Cheryl Jia Yi Neo, Siu Ling Wong, Kimberly A. Kline

## Abstract

*Enterococcus faecalis* is an opportunistic pathogen that persists in biofilm-associated infections despite robust neutrophil recruitment. How *E. faecalis* evades neutrophils remains poorly understood. Here, we investigated how *E. faecalis* biofilms alter effector functions in human neutrophils and a murine wound infection model. We found that *E. faecalis* biofilm suppresses neutrophil extracellular trap (NET) formation, bacterial engulfment, and neutrophil-mediated control of biofilm growth. This suppression is partly driven by production of lactic acid through the lactate dehydrogenases (LDH) of *E. faecalis*, which acidifies the extracellular environment, lowers neutrophil intracellular pH, and disrupts glycolysis, the tricarboxylic acid cycle, and the oxidative pentose phosphate pathway. In a murine wound infection model, loss of LDH increased neutrophil recruitment and NETosis. We also found that high bacterial density, a defining feature of biofilms, suppresses NETosis through a lactic acid-independent mechanism. Collectively, these findings identify biofilm immune evasion strategies mediated by bacterial metabolism and the high-density structural organization of biofilm-associated bacteria.

**Teaser:** *Enterococcus faecalis* biofilms suppress neutrophil defenses through lactic acid and high bacterial density.

## INTRODUCTION

*Enterococcus faecalis* is a gastrointestinal commensal and an opportunistic pathogen responsible for biofilm-associated infections, including infective endocarditis (IE), chronic wound infections, and catheter-associated urinary tract infections (CAUTI) (*1*, *2*). A key feature underlying *E. faecalis* pathogenesis is its ability to evade immune responses (*3*). Across different infection niches, *E. faecalis* persists despite neutrophil and macrophage recruitment. In infected wounds, *E. faecalis* induces neutrophil and macrophage infiltration, yet bacterial burdens remain detectable for days (*4*). Similarly, in IE, dense neutrophil infiltration and extensive NETosis within vegetations fail to eradicate biofilm microcolonies of *E. faecalis* (*5*). During CAUTI, catheter implantation triggers strong neutrophil influx and cytokine production, but *E. faecalis* efficiently persists on the catheter surface and urinary tract despite inflammation (*6*, *7*). These observations suggest that *E. faecalis* persistence is not due to insufficient innate immune infiltration, but rather due to active mechanisms that dampen or subvert innate immune effector functions.

*E. faecalis* interferes with both macrophage and neutrophil functions through a strategy that combines extracellular-driven immunosuppression with intracellular resistance to killing (*3*, *8*). During wound infection, *E. faecalis* can survive intracellularly within both immune and non-immune cells (*9*, *10*). Within keratinocytes and macrophages, a subpopulation of internalized bacteria survives and replicates in endosomal compartments that fail to mature and fuse with lysosomes (*9*). The intracellular lifestyle of *E. faecalis* in macrophages is regulated through the quorum sensing-regulated secreted protease gelatinase (GelE), which controls the transition between intracellular replication and extracellular dissemination (*11*). Intracellular bacterial clusters are also detected *in vivo* within neutrophils during acute wound infection. *In vitro*, phagocytosed *E. faecalis* suppresses rather than induces neutrophil death for up to 24 h and internalized bacteria persist and replicate within neutrophils (*10*). Moreover, *E. faecalis* strains expressing aggregation substance resist killing by both neutrophils and macrophages (*12*, *13*). Besides resisting intracellular killing, *E. faecalis* also dampens inflammation. *E. faecalis* inhibits LPS- or LTA-induced NF-κB activation in RAW264.7 macrophages and reduces NF-κB-dependent cytokine and chemokine production (*14*). During polymicrobial infection, *E. faecalis* dampens *Escherichia coli*-driven immune activation both *in vitro* and in a mouse model of CAUTI, where coinfection results in a blunted macrophage transcriptional response in the bladder and enhanced *E. coli* virulence (*14*). Mechanistically, *E. faecalis* suppresses macrophage immune responses through lactic-acid-mediated acidification of the extracellular environment (*15*). Lactic acid interaction with the lactate transporter MCT-1 and the receptor GPR81 leads to altered intracellular signaling leading to NF-κB inhibition. In a wound infection model, lactic acid-driven immunosuppression promotes prolonged *E. faecalis* persistence and enhances the fitness of co-infecting *E. coli* (*15*). *E. faecalis* also actively modulates neutrophil effector functions, including neutrophil extracellular trap (NET) release or NETosis, a response in which chromatin decorated with histones and antimicrobial proteins is released extracellularly to contain and kill microbes (*16*). *E. faecalis* alone fails to induce NETosis and attenuates NETosis triggered by *Staphylococcus aureus* during co-infection (*17*), indicating suppression of neutrophil antimicrobial mechanisms. However, the molecular mechanisms underlying neutrophil immunosuppression by *E. faecalis* remain undefined.

Previous studies of *E. faecalis*-neutrophil interactions have relied on planktonic bacteria (*10*, *12*, *17*), yet in the mammalian host this pathogen predominantly exists within biofilms (*2*). Globally, an estimated ∼69% of clinical *E. faecalis* isolates are biofilm-forming, with the highest prevalence (∼80%) among urinary isolates (*18*) (*2*). *In vivo* models further confirm that *E. faecalis* forms biofilms in the host, such as throughout the lower gastrointestinal tract of colonized mice (*19*), on catheters implanted in the mouse bladder (*20*), and on heart valve vegetations of a rat IE model (*5*). The extracellular polymeric substances found in *E. faecalis* matrix are eDNA (*21*), polysaccharides, such as the enterococcal polysaccharide antigen and β-1,6-linked poly-*N*-acetylglucosamine-containing exopolymers, (*22–24*), and proteins, such as the enterococcal surface protein (*25*, *26*). Biofilm growth profoundly reshapes the interactions between immune cells and bacteria by imposing structural barriers, increasing local bacterial density, and modifying its surrounding milieu with bacteria-secreted factors(*27–29*). For instance, *E. faecalis* biofilm produces L-lactate to acidify its microenvironment and inhibit *Pseudomonas aeruginosa* and *Klebsiella pneumoniae* growth (*29*, *30*). How such biofilm-associated changes in *E. faecalis* also alter immune cell functions remains less well understood. Although *E. faecalis* biofilm elicits a different response in macrophages and dendritic cells compared to planktonic bacteria (*31*), whether and how the biofilm-associated niche alters neutrophil effector functions is not known. Here, we demonstrate that *E. faecalis* biofilm actively suppresses NETosis and phagocytic uptake by human neutrophils. This suppression is mediated in part by *E. faecalis* lactic acid production, which suppresses ATP- and NADPH-producing metabolic pathways in neutrophils. In parallel, high bacterial density alone, which is an essential trait of biofilms, can also suppress NETosis in a lactic-acid independent manner. *In vivo*, *E. faecalis* LDH is associated with reduced neutrophil recruitment and diminished NET formation, promoting bacterial persistence. Together, these findings identify lactic acid production and biofilm density as potent immune evasion mechanisms that disarm neutrophils.

## RESULTS

### *E. faecalis* inhibits PMA-induced NETosis in human neutrophils

To investigate neutrophil-biofilm interactions in enterococcal infections, we employed a biofilm-neutrophil co-culture model in which human neutrophils were incubated with pre-formed biofilm of *E. faecalis* OG1RF (WT) for 4 hours (h) (**Fig. 1A**). We assessed NETosis by fluorescence microscopy using established markers (*32*) that distinguish early chromatin decondensation from later NET release. Histone H3 citrullination (H3Cit) and decondensed nuclei were used as early markers of NETosis, whereas NET extrusion and MPO-DNA co-localization were used to identify later-stage NET formation and release (**Fig. S1**). Similar to planktonic infection (*17*), neutrophils exposed to *E. faecalis* biofilm showed no evidence of NETosis compared to uninfected cells (**Fig. 1B-D**). Given our previous finding that *E. faecalis* partially suppresses *S. aureus*-induced NETosis (*17*), we also asked whether *E. faecalis* biofilm broadly suppresses NETosis triggered by other stimuli. We incubated biofilm with neutrophils in the presence of either phorbol 12-myristate 13-acetate (PMA) or ionomycin (IM), well-characterized inducers of NADPH oxidase (NOX)-dependent and -independent NETosis, respectively (*33*). *E. faecalis* inhibited PMA-induced NETosis, as evidenced by the total absence of H3Cit^+^ nuclei, decondensed nuclei, and NETosis (**Fig. 1B-D**). IM-induced NETosis hallmarks were detected in the presence of biofilm, including widespread histone H3 citrullination and ∼10% of neutrophils undergoing NETosis with released NETs co-localizing with MPO (**Fig. 1B-D**). IM-induced NETosis hallmarks were observed even after a 2-h pre-incubation of neutrophils with biofilm **(Fig. S2B-D**), showing that it is not suppressed by *E. faecalis*. Collectively, these results revealed that *E. faecalis* actively suppresses PMA-induced NETosis.

**Figure 1.**
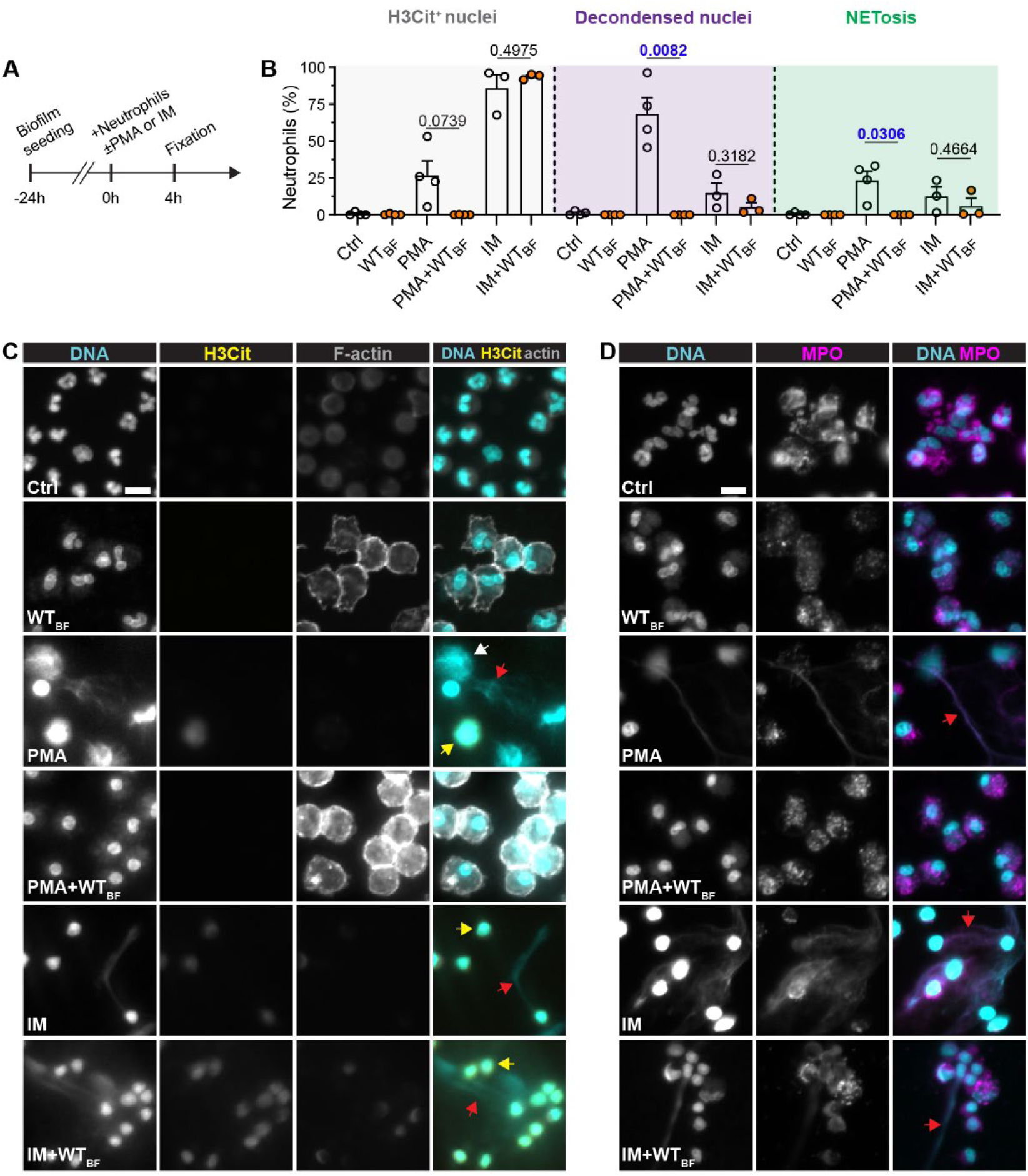
*E. faecalis* inhibits PMA-induced NETosis in human neutrophils. **A.** Experimental timeline for panels B-D. *E. faecalis* OG1RF WT biofilm (WT_BF_) was grown for 24 h, followed by simultaneous addition of human neutrophils and PMA (100 nM) or ionomycin (IM; 4 µM) at 0 h, and fixation at 4 h. **B.** Quantification of NETosis hallmarks from immunofluorescence microscopy images. Mean (%) citrullinated histone H3 stained nuclei (H3Cit^+^), decondensed nuclei, and NETs release (NETosis) from N = 3 - 4 independent experiments are shown, error = SEM. Statistical significance was assessed with two-tailed paired t-test. α = 0.05 with significant values shown in blue. **C-D.** Representative immunofluorescence microscopy images of neutrophils stained with DAPI, phalloidin actin staining, and either anti-H3Cit or anti-myeloperoxidase (MPO) antibodies. H3Cit images are from N = 3 - 4 and MPO images from N = 1. Scale = 10 µm, yellow arrow = H3Cit^+^ nuclei, white arrow = decondensed nuclei, red arrow = NETs; Ctrl = uninfected & unstimulated neutrophils in media.

### *E. faecalis* inhibits PMA-induced NETosis via lactic acid-mediated extracellular acidification

As the majority of infected neutrophils contained intracellular bacteria (**Fig. S2A**), we next tested whether internalized bacteria were suppressing PMA-induced NETosis. Despite inhibition of phagocytosis with cytochalasin D, the extracellular presence of *E. faecalis* biofilm continued to suppress NETosis (**Fig. S3A-B**). Thus, these findings demonstrate that *E. faecalis* inhibits NETosis extracellularly. Building on our earlier work identifying *E. faecalis*-derived lactic acid as an immunosuppressive factor in macrophages (*15*), we hypothesized that extracellular acidification could play a similar role in suppressing NETosis. To investigate this, we used an Δ*ldh1*Δ*ldh2* deletion mutant (Δ*ldh*) deriving from the OG1RF strain, which lacks the two lactate dehydrogenases that convert pyruvate to lactic acid. Because lactic acid rapidly dissociates into lactate and H^+^ (*34*), deletion of *ldh1* and *ldh2* abolishes this major driver of extracellular acidification (*35*). We harvested and measured the pH of 4-h biofilm supernatants from the WT strain (pH ∼4.8) and the Δ*ldh* mutant (pH ∼7.0) and proceeded to evaluate their effect on PMA-induced NETosis (**Fig. 2C**). WT supernatant completely inhibited the NETosis process, whereas the inhibitory effect was only partial using Δ*ldh* supernatant, with neutrophils displaying nuclear chromatin decondensation (**Fig. 2A & Fig. S3C**). Direct acidification of Δ*ldh* supernatant with lactic acid to achieve pH∼4.8 was sufficient to decrease decondensation in the nucleus and inhibit PMA-induced NETosis in the absence of bacteria (**Fig. 2B & Fig. S3D**). Similarly, gradual acidification of neutrophils with HCl alone inhibited PMA-induced chromatin decondensation in the nucleus (**Fig. 2D & Fig. S3E**). Neutrophils maintained comparable viability to the media control under all conditions tested, ruling out absence of NETosis because of cell death, as shown by the preservation of the multilobular nucleus morphology (**Fig. S3C-E**) and cell death quantification with propidium iodide (**Fig. S3F**). To investigate whether NETosis inhibition is specific to the OG1RF strain, we tested other clinical isolates of *E. faecalis* encoding different virulence factors (**Table S1**). All strains tested inhibited PMA-induced NETosis, apart from strain TTSHW-EF37, which exhibited reduced inhibition of histone H3 citrullination (**Fig. S4A-C**) and correlated with reduced media acidification because of impaired growth (**Fig. S4D**). To distinguish the contribution of lactate as a signaling molecule from acidification by H⁺, we also titrated neutrophils with either sodium L- or D-lactate across a concentration range spanning physiological and pathological levels reported in humans (*36*). Neither L-lactate nor D-lactate alone induced NETosis (**Fig. 2E, F & Fig. S5A, B**) and both exhibited a trend towards reducing NETosis hallmarks. Taken together, we conclude that inhibition of PMA-induced NETosis by *E. faecalis* is primarily driven by lactic acid-mediated acidification, which inhibits chromatin decondensation in the nucleus.

**Figure 2.**
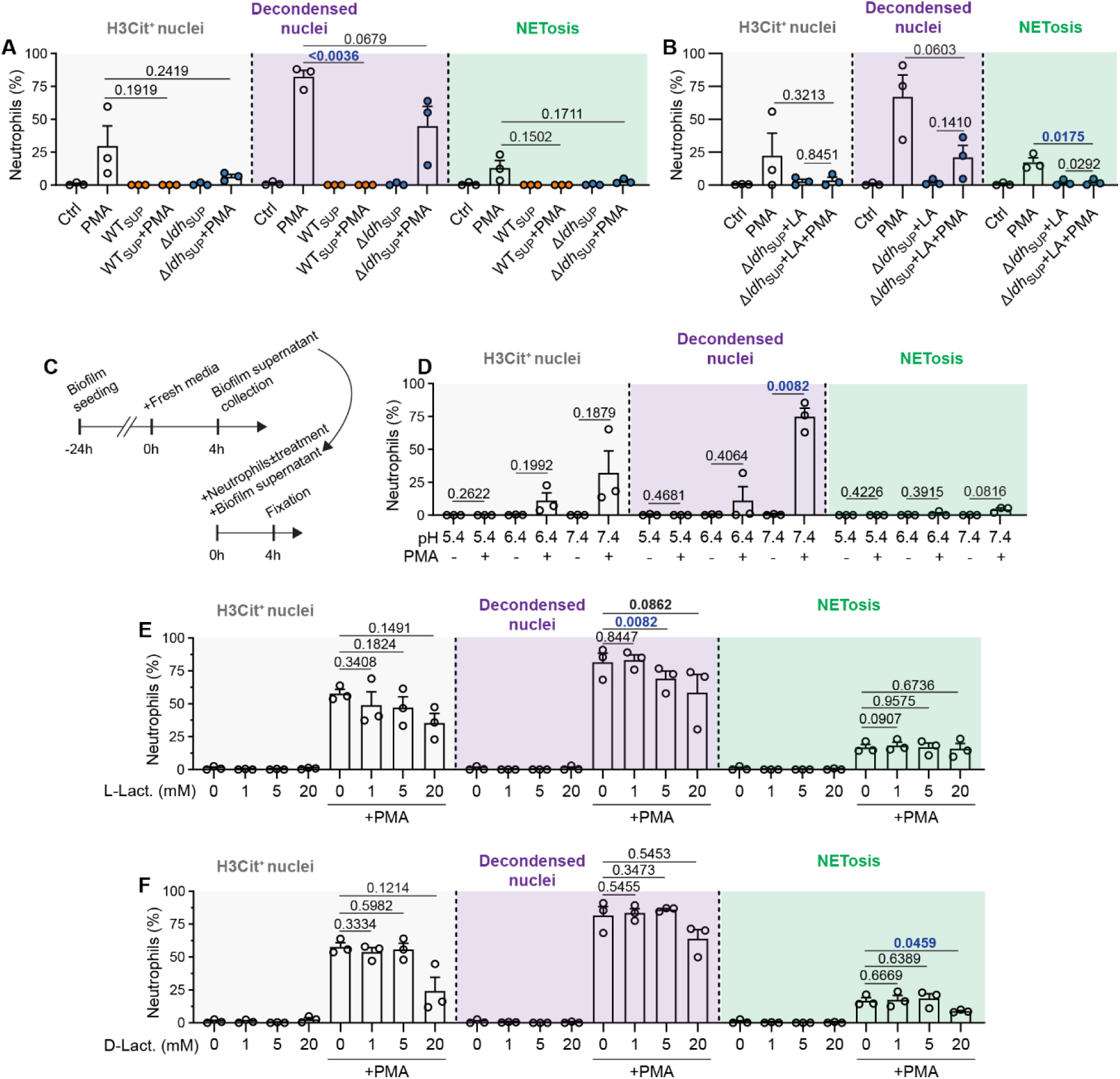
*E. faecalis* inhibits PMA-induced NETosis via lactic acid-mediated extracellular acidification. **A-B &D-F.** Quantification of NETosis hallmarks from immunofluorescence microscopy images. Human neutrophils were incubated for 4 h in WT or Δ*ldh* biofilm supernatant (WT_SUP_ & Δ*ldh*_SUP_ respectively) in (A), in acidified Δ*ldh* biofilm supernatant with LA in (B), in acidified media with HCl in (D), in media with L-lactate in (E), or D-lactate in (F), with or without co-stimulation with PMA added at 0 h. Mean (%) citrullinated histone H3 stained nuclei (H3Cit^+^), decondensed nuclei, and NETs release (NETosis) from N = 3 independent experiments are shown, error = SEM. Panels E and F were generated from the same experiment and therefore share the same control and PMA samples; the data are presented in separate graphs for clarity. Statistical significance was assessed with two-tailed paired t-test. α = 0.05 with significant values shown in blue. **C.** Experimental timeline for B-C. WT and Δ*ldh* biofilm was grown for 24 h, followed by exchange of old media with fresh media and an additional 4-h incubation. At 4h, biofilm supernatant was harvested and added to human neutrophils with or without PMA (100 nM) or lactic acid (LA; 20mM) at 0 h and fixed at 4 h.

### Acute acidification by lactic acid impairs *E. faecalis* engulfment by neutrophils

Since *E. faecalis*-derived lactic acid inhibits NETosis, we asked whether this inhibition extends to phagocytosis. First, we compared the ability of neutrophils to engulf bacteria from the WT and Δ*ldh* biofilm. Nearly all neutrophils exposed to pre-formed biofilms contained dense intracellular bacterial clusters by 1 h post-incubation, regardless of the presence of *ldh1* and *ldh2* (**Fig. 3A**). However, whether engulfment resulted in killing remained unclear. To assess whether *ldh1* and *ldh2* affect bacterial susceptibility to neutrophil killing, we grew WT and Δ*ldh* biofilms for 24 h. Spent media was then replaced with fresh media, and baseline biofilm CFU was quantified, which was comparable between the 2 strains (**Fig. 3B**, orange & blue dashed line). We incubated biofilms for an additional 4 h in the presence or absence of neutrophils. Δ*ldh* biofilm showed reduced growth compared with WT biofilm by 4 h post-incubation in the absence of neutrophils (**Fig. 3B**). In the presence of neutrophils, WT biofilm grew equally well, while *Δldh* biofilm growth was further impaired (**Fig. 3B**). This growth impairment, however, was independent of bacterial engulfment, since cytochalasin D had no effect on the ability of neutrophils to restrict *Δldh* biofilm growth (**Fig. 3B**).

**Fig. 3.**
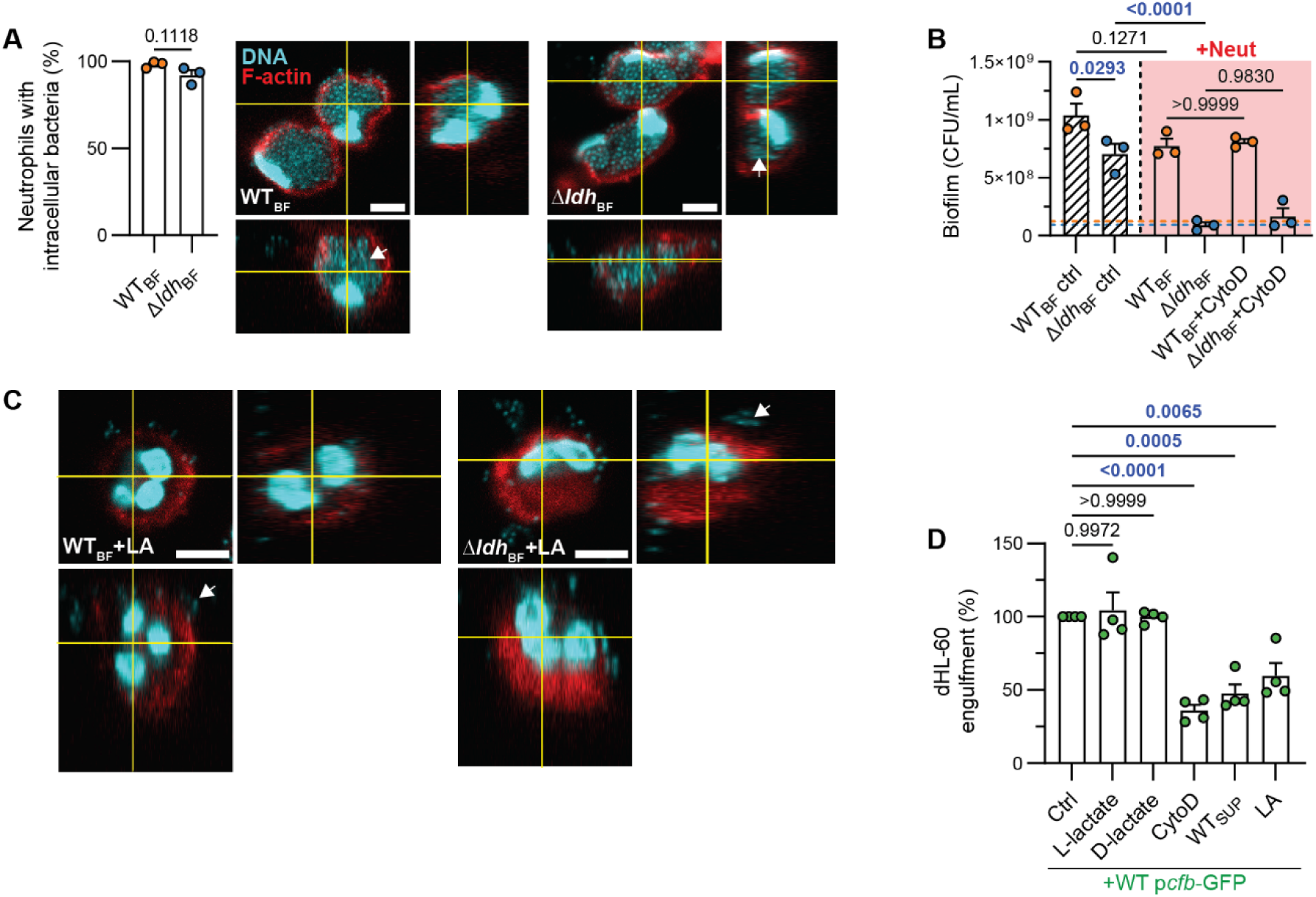
Acute acidification by lactic acid impairs *E. faecalis* engulfment by neutrophils. **A.** Mean (%) of human neutrophils with intracellular bacteria after a 1-h incubation with pre-formed WT and Δ*ldh* biofilm (WT_BF_ and Δ*ldh*_BF_ respectively). A representative Z-stack slice and respective orthogonal views are shown. White arrows indicate the dense intracellular bacterial clusters. N = 3, error = SEM, scale = 5 µm. Statistical significance was assessed with t-test. **B.** Mean CFU/mL of *E. faecalis* WT and Δ*ldh* biofilm at 4 h in the absence or presence of human neutrophils (Neut) and cytochalasin D (30uM; cytoD). Orange and blue dashed lines correspond to CFU at 0 h for WT and Δ*ldh* biofilm respectively. N = 3, error = SEM. Statistical significance was assessed with one-way ANOVA followed by Tukey’s multiple comparisons test. N = 3, error = SEM. **C**. Representative Z-stack slice and respective orthogonal views of neutrophils incubated with either WT or Δ*ldh* biofilm, and lactic acid (20 mM; LA) at 1 h. White arrows indicate bacteria attached to neutrophils extracellularly. N = 3, scale = 5 µm. **D.** Normalized engulfment of *E. faecalis* WT P*c@*-GFP by dHL-60 cells at 1 h quantified by flow cytometry. N = 4, error = SEM. Statistical significance was assessed with one-way ANOVA followed by Tukey’s multiple comparisons test; α = 0.05 with significant values shown in blue.

Following addition of fresh media at 0 h, acidification in the biofilm-neutrophil co-culture model develops gradually over the course of 4 h. By contrast, neutrophils encountering biofilm in the host are likely exposed to an already acidified microenvironment adjacent to the mature biofilm. To model this pre-existing acidification, we added lactic acid and neutrophils to pre-formed biofilm at 0 h and quantified bacterial engulfment at 1 h. Lactic acid markedly reduced engulfment of bacteria from both WT and Δ*ldh* biofilms (**Fig. 3C**), indicating that immediate acidification impairs neutrophil phagocytosis. To quantify this observation, we incubated differentiated HL-60 (dHL-60) cells, which provide a tractable neutrophil-like model, with WT OG1RF biofilm constitutively expressing GFP from a plasmid (WT P*c@*-GFP) and quantified engulfment by flow cytometry. Addition of lactic acid or WT biofilm supernatant significantly reduced phagocytosis to levels comparable to cytochalasin D treatment (**Fig. 3D**), whereas L- or D-lactate alone had no effect. These results show that acute lactic acid-mediated acidification impairs neutrophil engulfment of biofilm-associated bacteria. Thus, lactic acid production enables *E. faecalis* to evade neutrophil phagocytosis.

### *E. faecalis-*derived lactic acid suppresses glycolysis, the TCA cycle, and the oxidative PPP (oxPPP) in neutrophils

Effector functions in neutrophils require ATP generation to meet energy demands (*37*). Given the inhibition of PMA-induced NETosis, reduced bactericidal activity, and impaired bacterial engulfment in the presence of *E. faecalis*-derived lactic acid, we investigated whether extracellular acidification affects neutrophil metabolism. We incubated neutrophils with WT or Δ*ldh* biofilm supernatant and measured heat production using microcalorimetry as an indicator of their metabolic activity. Heat output was lower in the presence of WT supernatant compared to fresh media, while Δ*ldh* supernatant had no significant effect (**Fig. 4A, B**). Microcalorimetry using dHL-60 showed the same effect, with WT supernatant consistently suppressing heat production (**Fig. S6A, B**). Together, these data suggest that *E. faecalis* lactic acid suppresses neutrophil metabolism.

**Fig. 4.**
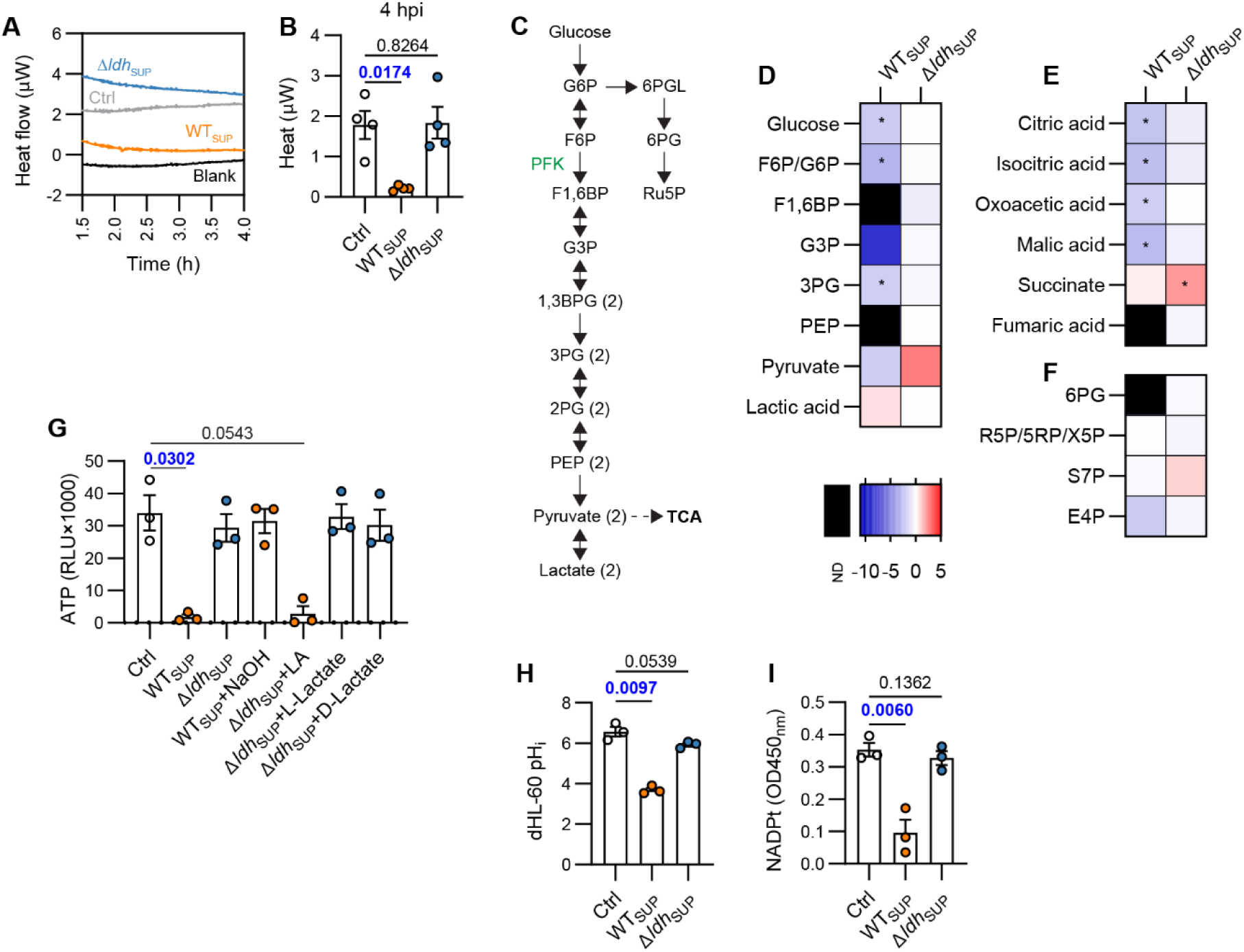
*E. faecalis-*derived lactic acid suppresses glycolysis, the TCA cycle, and the oxPPP in neutrophils. **A.** Representative thermogram of human neutrophils treated with fresh media (Ctrl), or biofilm supernatant from WT (WT_SUP_) or Δ*ldh* (Δ*ldh*_SUP_) for 4 h. Media alone (blank) was included. Heat measurements start after 1.5 h due to the calScreener initial calibration phase. N = 4. **B.** Mean heat produced by neutrophils at 4 h post-incubation. N = 4, error = SEM. Statistical significance was assessed with two-tailed paired t-test. **C.** Schematic representation of glycolysis and part of oxPPP including metabolites analyzed with LC-MS. Metabolites labeled with (2) are generated as two molecules per molecule of glucose. Single arrows indicate irreversible steps, double arrows indicate reversible steps, and dashed arrows indicate pathway branches of TCA not shown in full. Key enzymes are shown in green. **D-F.** LC-MS heatmaps showing log2 fold changes in metabolite abundance in neutrophils after exposure to WT_SUP_ or Δ*ldh*_SUP_ for 4 h relative to untreated control cells. Metabolites from glycolysis (D), the TCA cycle (E), and the PPP (F) are shown. N = 5, color scale indicates relative metabolite abundance, with red denoting increased and blue decreased levels relative to control, and black denoting not detected (ND) metabolites. Statistics were calculated on the biological replicate mean peak areas using a two-tailed Welch’s t-test to compare WT_SUP_ with media and Δ*ldh*_SUP_ with media for each metabolite. * = P < 0.05, ** = P < 0.01, *** = P < 0.001, **** = P < 0.0001. **G.** ATP levels in neutrophils after incubation with media alone (Ctrl), WT_SUP_ or Δ*ldh*_SUP_ for 4 h. NaOH, lactic acid (LA), sodium L- or D-Lactate were added where indicated. Data is presented as relative luminescence units (RLU), N = 3, error = SEM. Statistical significance was assessed with two-tailed paired t-test. **H.** Mean intracellular pH (pH_i_) in dHL-60 cells after incubation with media alone (Ctrl), WT_SUP_ or Δ*ldh*_SUP_ for 4 h. N = 3, error = SEM. Statistical significance was assessed with two-tailed paired t-test. **I**. Total NADP^+^/NADPH (NADPt) in neutrophils incubated with media alone (Ctrl), WT_SUP_ or Δ*ldh*_SUP_ for 4 h. N = 3, error = SEM. Statistical significance was assessed with two-tailed paired t-test.; α = 0.05 with significant values shown in blue.

To identify which metabolic pathways are affected, we performed LC-MS on neutrophil cell lysates following exposure to WT or Δ*ldh* biofilm supernatant. WT supernatant caused a broad suppression of glycolytic intermediates compared to control cells, with decreased glucose, glucose-6-phosphate/fructose-6-phosphate (G6P/F6P), glyceraldehyde-3-phosphate (G3P; undetectable in 4 out of 5 replicates), and 3-phosphoglycerate (3PG), and undetectable fructose-1,6-bisphosphate (F1,6BP) and phosphoenolpyruvate (PEP), indicating strong depletion (**Fig. 4C, D, Supplementary Data File 1)**. By contrast, treatment with Δ*ldh* biofilm supernatant maintained glycolysis in neutrophils, with increased pyruvate levels relative to the control (**Fig. 4C, D, Supplementary Data File 1)**. WT biofilm supernatant had a similar effect on dHL-60 glycolysis, which was largely reversed in the absence of LDH for most metabolites apart from F6P/G6P (**Fig. S6D, Supplementary Data File 2**). Although mature neutrophils rely primarily on glycolysis and contain relatively few mitochondria, mitochondrial ATP can contribute to neutrophil activation (*38*). We therefore assessed whether bacterial supernatants altered mitochondrial metabolism by quantifying TCA cycle intermediates (**Supplementary Data File 1**). WT biofilm supernatant resulted in the decrease of citrate, isocitrate, and oxoacetate and marked depletion of fumarate (**Fig. 4E**). By contrast, Δ*ldh* supernatant largely preserved these metabolites, with succinate increased relative to the control. dHL-60 also exhibited a marked decrease in TCA metabolic intermediates in the presence of WT biofilm supernatant, which partially was maintained even in Δ*ldh* supernatant (**Fig. S6E, Supplementary Data File 2**). Consistent with the disruption of these central ATP-generating pathways, WT supernatant significantly reduced intracellular ATP levels in neutrophils, with a similar trend observed in dHL-60 cells (**Fig. 4G** & **Fig. S6C**). Neutralized WT supernatant and sodium lactate-supplemented Δ*ldh* supernatant did not reduce ATP, whereas lactic acid supplementation caused a sharp decrease (**Fig. 4G**). Together, these results show that *E. faecalis* lactic acid-mediated acidification suppresses glycolysis and the TCA cycle, leading to ATP depletion in neutrophil and dHL-60 cells.

The activity of phosphofructokinase (PFK), the rate-limiting enzyme that converts F6P to F1,6BP during glycolysis (**Fig. 4C**), decreases in low pH (*39–41*), suggesting a potential mechanism for the glycolysis decrease. Consistent with this, the intracellular pH was significantly lower in dHL-60 cells exposed to WT biofilm supernatant, whereas Δ*ldh* biofilm supernatant had no effect (**Fig. 4H**), indicating that lactic-acid mediated intracellular acidification may suppress glycolysis at the level of PFK and explain why F1,6BP was undetected. Because L-lactate can be transported together with protons through monocarboxylate transporters such as MCT-1 and MCT-4 (*42*), we next asked whether MCT-dependent lactate transport contributed to NETosis inhibition. However, MCT inhibition alone partially reduced PMA-induced NETosis in neutrophils and combined treatment with MCT inhibitors and WT biofilm supernatant further reduced NETosis (**Fig. S7**), suggesting that intracellular acidification occurs via other mechanisms.

PMA-induced NETosis relies on NOX2-generated ROS, which require NADPH as an essential electron donor (*43*). The oxPPP produces NADPH, in which glucose-6-phosphate dehydrogenase (G6PDH) converts G6P to 6-phosphogluconolactone (6PGL) and 6-phosphogluconate dehydrogenase (6PGDH) converts 6-phosphogluconate (6PG) to ribulose-5-phosphate (Ru5P), with each step generating NADPH (**Fig. 4C**) (*37*). Since *E. faecalis* suppresses PMA-induced NETosis, we investigated whether this involves disruption of the oxPPP. 6PG was depleted in both neutrophils and dHL-60 cells exposed to WT supernatant, indicating suppression of this pathway (**Fig. 4F & Fig. S6F, Supplementary Data 1 & 2**). In primary neutrophils, 6PG was retained in the presence of Δ*ldh* supernatant, indicating lactic acid–dependent suppression of the oxPPP. In dHL-60 cells, 6PG was depleted by both WT and Δ*ldh* supernatants, indicating a lactic acid– independent depletion. Consistent with the undetectable levels of 6PG in the presence of WT biofilm supernatant in neutrophils, their total NADP^+^/NADPH levels were also decreased (**Fig. 4I**). With the exception of decreased erythrose-4-phosphate (E4P) levels in neutrophils, metabolites of the non-oxidative PPP branch were similar across conditions in both neutrophils and dHL-60 (**Fig. 4F & Fig. S6F**). These data show that *E. faecalis* suppresses the oxPPP in a lactic acid-dependent manner, which in turn limits NADPH availability. Since NADPH is required for NOX2 activity, this result provides a plausible explanation for the PMA-induced NETosis suppression in the presence of WT biofilm supernatant.

### Δ*ldh* biofilm suppresses NETosis in a density-dependent manner

Given the importance of extracellular acidification in inhibiting NETosis, and the observation that PMA-induced NETosis was restored when neutrophils were incubated with Δ*ldh* biofilm supernatant, we next examined whether neutrophils exposed to intact Δ*ldh* biofilm were able to undergo NETosis. Neutrophils failed to undergo NETosis in the presence of Δ*ldh* biofilm, with or without PMA stimulation, suggesting that additional bacterial mechanisms inhibit NETosis independently of acidification. (**Fig. 5A & Fig. S8**). Addition of sodium L- or D-lactate to Δ*ldh* biofilm also did not trigger NETosis (**Fig. S9A, B**). As the majority of neutrophils internalized Δ*ldh* bacteria (**Fig. 3A**), we further assessed whether internalized bacteria could inhibit PMA-induced NETosis. However, preventing bacterial uptake with cytochalasin D did not impact the ability of Δ*ldh* biofilm to suppress NETosis (**Fig. 5A & Fig. S8**), indicating the inhibition of NETosis via additional extracellular contact-dependent mechanisms in *E. faecalis*. To determine whether this inhibition was mediated by the biofilm ultrastructure alone or required active contact-dependent bacterial processes, we tested PFA-fixed Δ*ldh* biofilms, in which bacterial activity is abolished while biofilm ultrastructure is preserved. Neither the fixed Δ*ldh* biofilm alone nor supplementing with PMA triggered NETosis (**Fig. S10A, B**), suggesting that structural components may mediate this inhibition. To determine whether this inhibition depended on intact biofilm ultrastructure, we mechanically dispersed PFA-fixed Δ*ldh* biofilms by scraping and assessed whether the disrupted biofilm material retained the ability to suppress NETosis. NETosis suppression persisted after biofilm disruption, regardless of the presence of PMA (**Fig. S10A, B**). Biofilm inherently presents a high multiplicity of infection (MOI) to incoming neutrophils. Thus, to determine whether NETosis suppression is bacterial density-dependent, we infected neutrophils with stationary phase planktonic Δ*ldh* at increasing MOI, including MOI 1000 which is equivalent to the MOI in the biofilm infection assays. MOI 1000 inhibited all hallmarks of NETosis in the presence of PMA (**Fig. 5B, C**). By contrast, neutrophils infected with MOI 100 and 10, exhibited decondensed nuclei and NETosis in the presence of PMA (**Fig. 5B, C**). Thus, Δ*ldh* biofilm inhibits NETosis in a density-dependent manner and not via its ultrastructure or a biofilm-specific extracellular component. Together, these data show that, in addition to lactic acid, *E. faecalis* can inhibit PMA-induced NETosis potentially through structural bacterial components shared by planktonic and biofilm bacteria, but only when bacteria are present at high density, as commonly occurs within biofilms.

**Figure 5.**
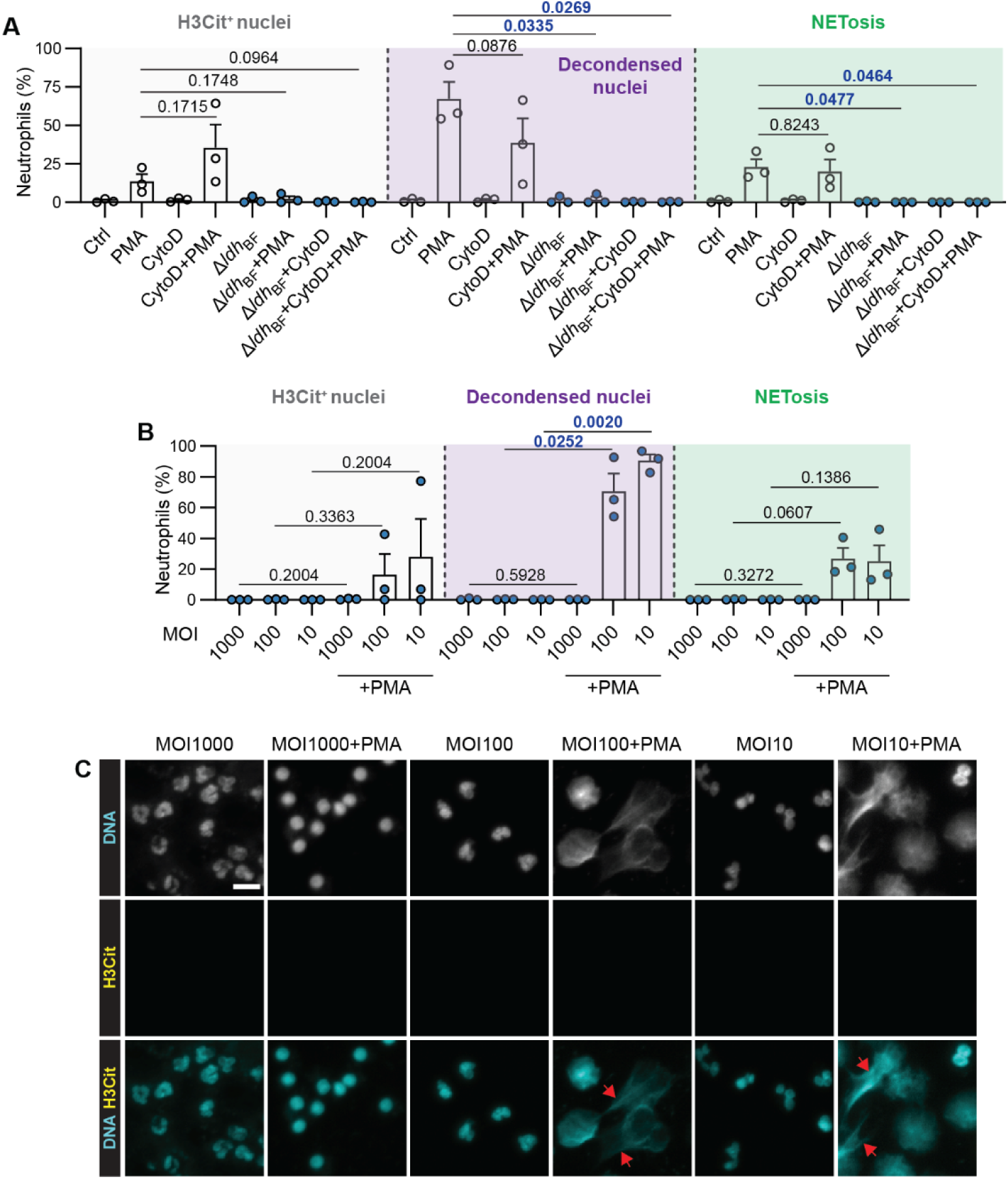
Δ*ldh* biofilm suppresses NETosis in a density-dependent manner. **A-B.** Quantification of NETosis hallmarks from immunofluorescence microscopy images. Mean (%) citrullinated histone H3 stained nuclei (H3Cit^+^), decondensed nuclei, and NETs release (NETosis) from N = 3 are shown, error = SEM. In 5A, human neutrophils were incubated with Δ*ldh* biofilm (Δ*ldh*_BF_) in the presence of cytochalasin D (cytoD; 30 μM) and PMA (100 nM) as indicated for 4 h. In 5B, human neutrophils were incubated with Δ*ldh* planktonic bacteria at MOI 1000, 100, or 10 with PMA as indicated for 4 h. Statistical significance was assessed with a two-tailed paired t-test.. α = 0.05 with significant values shown in blue. **C.** Representative images of neutrophils for 5B with nuclei stained for DAPI and citrullinated histone H3 (H3Cit). N = 3, scale = 10 µm, red arrows = NETs.

### *E. faecalis* LDH limits neutrophil recruitment and NET formation *in vivo*

We previously showed in a murine wound infection model that bacterial burdens of WT and Δ*ldh* strains are comparable at 1 day post-infection (dpi) but by 7 dpi, Δ*ldh* displays an approximately 10-fold reduction in bacterial load, suggesting enhanced clearance in the absence of lactate dehydrogenase (*15*). Because neutrophils are key mediators of bacterial clearance in acute wound infections (*4*), we next asked whether LDH influences neutrophil recruitment *in vivo*. We quantified neutrophils (CD45⁺ CD11b⁺ Ly6G⁺) in WT- and Δ*ldh*-infected wounds at 1 and 7 dpi. Consistent with the comparable CFU between WT and Δ*ldh*-infected wounds at 1 dpi (*15*), loss of LDH did not affect neutrophil recruitment or their proportion among immune cells at this time point (**Fig. 6A, C**) despite higher levels of CXCL1 and CCL2 chemokines at 1 dpi (**Fig. 6E, G**). Accordingly, by 7 dpi, WT-infected wounds exhibited decreased neutrophil infiltration compared to Δ*ldh*-infected wounds (**Fig. 6B, D**), coinciding with the impaired bacterial clearance previously observed at this time point (*15*). The proportion of neutrophils among total immune cells was likewise lower in WT infection, while CXCL1 and CCL2 levels were comparable by this time point (**Fig. 6F, H**). Next, to determine whether LDH also limits NET formation *in vivo*, we quantified H3Cit relative to total histone H3 in wound lysates. Although the H3Cit/H3 ratio was comparable at 1 dpi (**Fig. 6I**), it was higher in Δ*ldh*-infected wounds by 7 dpi (**Fig. 6J**), indicating enhanced NETosis in the absence of LDH. Together, these data suggest that LDH activity dampens early chemokine production, leading to reduced neutrophil recruitment and NET formation at later stages of infection, consistent with its suppressive effects on neutrophil function observed *in vitro*.

**Fig. 6.**
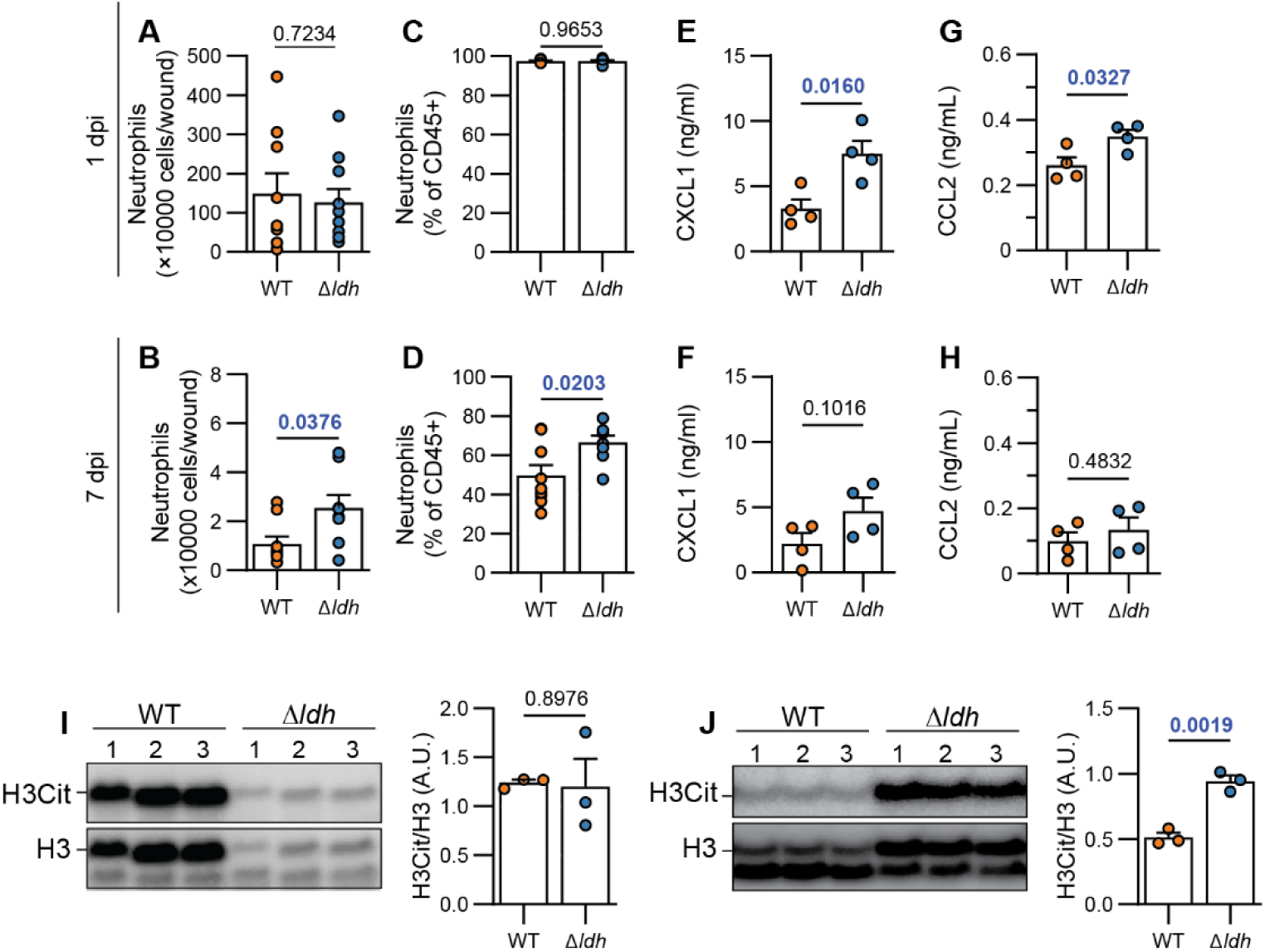
*E. faecalis* LDH limits neutrophil recruitment and NET formation *in vivo.* **A-D.** Mean absolute (A, B) and relative (C, D) neutrophil number in WT- and Δ*ldh* infected murine wounds at 1 and 7 dpi, assessed by flow cytometry. Samples derived from N = 2 independent experiments, n = 8 – 10 mice per condition, error = SEM. Statistical significance was assessed with unpaired t-test. **E-H.** Mean CXCL1 and CCL2 concentration from WT- and Δ*ldh* infected murine wounds at 1 and 7 dpi assessed with ELISA. N = 2, n = 4, error = SEM. Statistical significance was assessed with Welch’s t-test. **I, J.** Detection and quantification of citrullination of H3 histone (H3Cit) levels by western blot at 1 (I) and 7 (J) dpi in a WT- and Δ*ldh* infected murine wounds. Statistical significance was assessed with Welch’s t-test. N = 2, n = 3, error = SEM, A.U. = arbitrary units; α = 0.05 with significant values shown in blue.

## DISCUSSION

Our findings identify acidification as a mechanism by which *E. faecalis* biofilms evade NETosis and neutrophil-mediated bacterial engulfment (**Fig. 7A**). Bacterial lactic acid production acidifies the extracellular environment, lowers neutrophil intracellular pH, and suppresses central carbon metabolism, thereby reducing ATP and NADPH availability and impairing PMA-induced NETosis. Consistent with these *in vitro* findings, loss of bacterial LDH enhances neutrophil recruitment and NET formation in a murine wound infection model, linking *E. faecalis* metabolic activity to impaired innate immune clearance during infection (**Fig. 7B**). Moreover, we uncovered a second, lactic acid-independent contact-dependent mode of NETosis suppression, in which high-density biofilms create a suppressive microenvironment for NETosis (**Fig. 7A**). Together, these results reveal that the high density of *E. faecalis* within biofilms enables two modes of neutrophil suppression: (i) efficient acidification of the surrounding microenvironment through robust lactic acid production and (ii) a multitude of simultaneous interactions at the biofilm-neutrophil interface, which may amplify inhibitory signaling. Importantly, high cell density is an inherent property of the biofilm and the only state in which bacteria sustain such localized high densities in the host.

**Figure 7.**
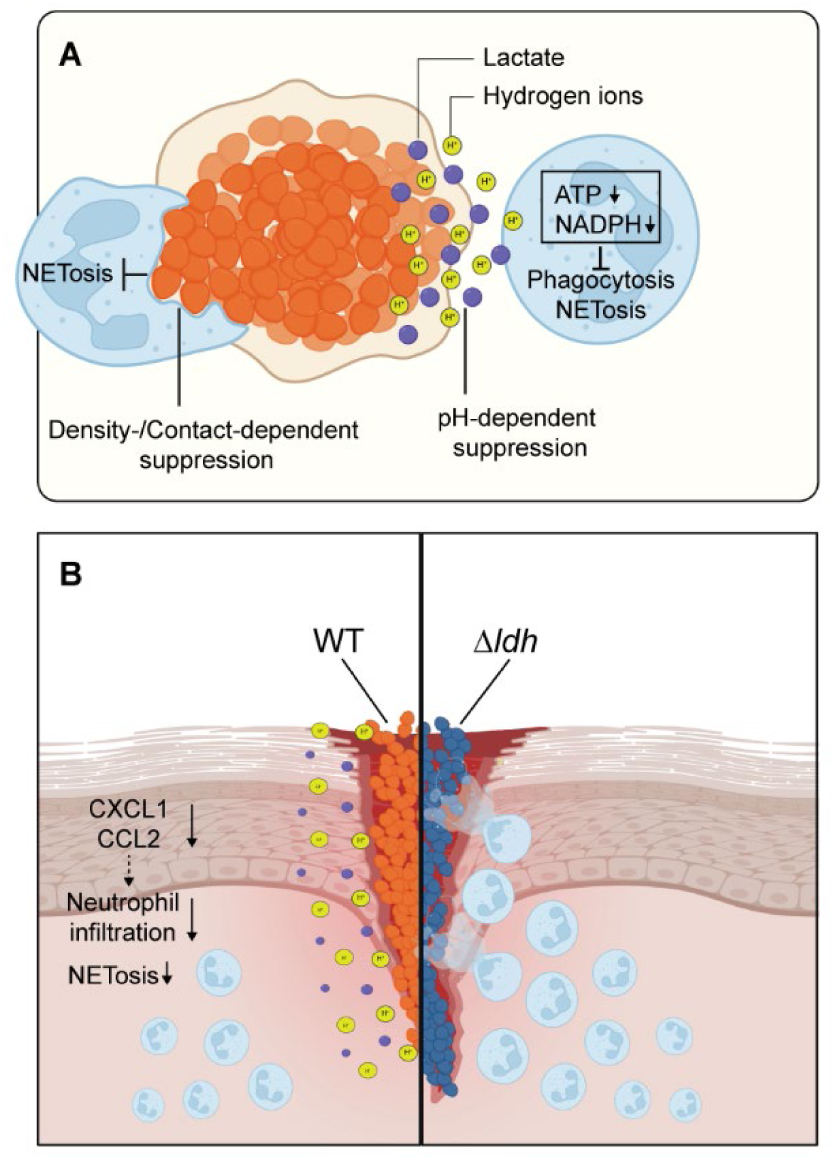
*E. faecalis*-derived lactic acid and density-dependent contact suppress neutrophil effector functions. **A.** WT *E. faecalis* biofilms produce lactic acid, resulting in extracellular acidification that lowers neutrophil intracellular pH and suppresses metabolic pathways (glycolysis, TCA, and oxPPP). This metabolic rewiring reduces NADPH and ATP availability and impairs NETosis and bacterial engulfment. In host microenvironments where acidification is not established despite lactic acid production, biofilm can still suppress NETosis through a supernatant-independent, density/contact-dependent mechanism, likely mediated by extracellular bacterial components. **B.** In a murine wound infection model, *E. faecalis* LDH suppresses chemokine secretion (CXCL1, CCL2) and limits neutrophil infiltration and NETosis. Created in BioRender. Antypas, H. (2026) https://BioRender.com/a9lpr6b

Inflammation is associated with a decrease in the extracellular pH of the local host microenvironment (*44–46*), and previous *in vitro* studies using HCl-acidified media have shown that extracellular acidification reduces ROS production, inhibits PMA-induced NETosis, delays apoptosis, impairs migration, and alters particle uptake by neutrophils (*47–51*). Our findings place these observations into a bacterial infection context by showing that extracellular acidification can be driven by a pathogen-derived metabolite during biofilm growth. This immune-evasion mechanism is likely to be broadly applicable to other lactic-acid–producing and acidifying bacteria. Mechanistically, we show that acidification by *E. faecalis*-derived lactic acid disrupts neutrophil central carbon metabolism, including glycolysis, the TCA cycle, and the oxPPP, thereby limiting ATP and NADPH availability, which are essential for NOX2-dependent NET formation. This is consistent with studies showing that NET formation requires glycolytic ATP production (*52–54*) and that PMA-induced NETosis depends on increased oxPPP activity to generate NADPH for NADPH oxidase-mediated superoxide production (*55*). This metabolic suppression may also explain our previous findings showing that *E. faecalis* reduces *S. aureus*-induced NETosis (*17*). Our findings are also in line with reports that acidified media reduce neutrophil glucose uptake and lactate production (*51*), and that lactic acid or acidic pH suppress glycolysis, ATP levels, and inflammatory cytokine production in mast cells and monocytes (*56*, *57*). By contrast, HCl-mediated acidification has been reported to enhance zymosan engulfment and preserve phagocytosis of opsonized non-viable *S. aureus* bioparticles (*47*, *51*), suggesting that the impaired engulfment of *E. faecalis* observed here may reflect distinct uptake mechanisms for live biofilm bacteria versus non-viable particles. Finally, because extracellular acidification reduced neutrophil intracellular pH, this is likely impairing the activity of glycolytic enzymes such as PFK (*39–41*), thereby limiting glycolytic flux. Such a mechanism would be consistent with our inability to detect F1,6BP, the product of the PFK-catalyzed reaction, and may explain the suppression of glycolysis and NET formation observed during *E. faecalis* infection. How *E. faecalis*-derived lactic acid decreases neutrophil intracellular pH remains to be investigated.

Δ*ldh* biofilms still suppress PMA-induced NETosis, despite the loss of lactic acid-mediated acidification. This suppression does not appear to require metabolically active bacteria, phagocytic uptake, or intact biofilm architecture, as it persisted even after PFA fixation and mechanical disruption of biofilm. Moreover, the ability of high MOI planktonic Δ*ldh* bacteria to similarly suppress PMA-induced NETosis argues against a mechanism that is linked to biofilm-specific metabolic changes. Instead, these findings suggest that *E. faecalis* can inhibit NETosis when neutrophils encounter a sufficiently high bacterial burden, likely through contact-dependent interactions with abundant bacterial surface-associated or structural components. Biofilms may therefore amplify this mechanism by creating a microenvironment in which neutrophils are exposed to extremely high bacterial densities. We also note that the MOI 1000 planktonic condition is an artificial *in vitro* approach with no *in vivo* planktonic counterpart. Thus, the biofilm is the only physiological state in which *E. faecalis* sustains such high densities in the host. The mechanism underlying the lactic acid-independent inhibition remains to be determined. We cannot exclude that *E. faecalis* may also suppress neutrophil metabolic activity in a contact-dependent manner, thus depriving neutrophils from ATP and NADPH required for NETosis. Impairment of oxPPP has been shown in neutrophils infected by *S. aureus*, which decreases the expression of G6PDH, associated with the upregulation of adenosine receptor 2A (*58*). Group B *Streptococcus* also suppresses the neutrophil oxidative burst and inhibits NETosis via its sialylated capsular polysaccharides interacting with Siglec-9 in neutrophils (*59*). Alternatively, *E. faecalis* contact-dependent inhibition may interfere with PMA-induced intracellular signalling cascades, including Raf-MEK-ERK, Akt, and p38 MAPK, which are activated during PMA-induced NETosis (*60*, *61*). Although *ldh1* and *ldh2* are conserved across *E. faecalis* strains, this secondary mechanism may be particularly relevant in well-buffered host environments, where extracellular acidification is limited or rapidly neutralized and lactic acid alone may be insufficient to suppress NETosis. Under these conditions, the high local bacterial burden characteristic of biofilm infections could provide an acidification-independent route to inhibit neutrophil activation.

Lactic acid can influence immune cell responses not only via H^+^-dependent acidification but also through lactate-dependent signaling (*62*). Lactate promotes neutrophil mobilization in the bone marrow (*63*) and decreases neutrophil apoptosis (*64*). Indirectly, gut microbiota-derived D-lactate also promotes neutrophil mobilization to liver by upregulating endothelium adhesion molecules for neutrophils (*65*). In our study, L- or D-lactate alone resulted only in a modest suppression of histone H3 citrullination at high concentrations (20 mM), showing a limited lactate-specific effect in our model compared to acidification. This differs from a previous report that 20 mM lactate induces NETosis in human neutrophils (*66*), which may be the result of neutrophil incubation in saline, unclear distinction between lactate and lactic acid use, and the classification of nuclear decondensation as NETosis without clear evidence of extracellular NET structures. Moreover, L-lactate produced inside the phagosomes of neutrophils by *S. aureus* can act as a metabolic danger signal, which when detected by mitochondria can trigger NETosis (*67*). However, we have not observed a comparable L-lactate-dependent induction of NETosis during *E. faecalis* infection, either in the current biofilm model or in our previous studies using planktonic bacteria at low MOI (*10*, *17*). In the biofilm setting, extracellular acidification may override or suppress any potential L-lactate-mediated NETosis. However, the absence of NETosis induction in low-MOI planktonic infection suggests that this pathway is not broadly activated by *E. faecalis*. Consistent with this, supplementation of Δ*ldh* biofilms with L- or D-lactate did not restore NETosis, further supporting the conclusion that *E. faecalis* employs additional acidification-independent strategies to suppress neutrophil responses.

Collectively, our findings identify the biofilm microenvironment itself as a determinant of immune evasion, integrating metabolic and contact-dependent mechanisms that suppress neutrophil effector functions. In the host, however, neutrophil responses are likely shaped not only by interactions with bacteria, but also by local physicochemical cues within the infection niche, nutrient availability, and sex-dependent differences. The overall phenotype therefore reflects the dominant signal within a given microenvironment. In our experimental system, extracellular acidification appears to be a major driver of the observed effects. However, its contribution is likely context dependent. The relatively static microenvironment of a wound bed may favor the local accumulation of lactic acid, thereby promoting acidification, whereas this effect may be attenuated in more dynamic and well-buffered environments, such as infective endocarditis vegetations. Nutrient availability within the infection niche may further modulate this mechanism. Glucose-rich environments such as diabetic wounds presumably could enhance *E. faecalis* LDH-driven acidification and neutrophil suppression, yet it has also been demonstrated that hyperglycemia simultaneously promotes host NETosis (*68*), making the net outcome hard to predict. Neutrophils restrict iron as an antimicrobial defense (*69*), yet iron restriction has been shown to increase *ldh1* expression and L-lactate production in *E. faecalis* (*29*). Host iron sequestration may therefore potentiate, rather than blunt, bacterial acidification. Similarly, the high local density of bacteria within a biofilm may favor the accumulation of metabolic products, steepen chemical gradients, and amplify contact-dependent interactions with neutrophils, unlike planktonic bacteria. Moreover, hormonal differences between sexes can influence immune responses (*70*). Future work should therefore focus on how bacterial lifestyle and metabolism, local host cues, and sex interact to determine how *E. faecalis* modulates immune responses in different infection settings, as these context-dependent mechanisms may require distinct therapeutic strategies.

## MATERIALS AND METHODS

### Bacterial cultures

Bacterial strains (**Table S1**) were routinely cultured in brain heart infusion (BHI) broth at 37°C without shaking, in ambient air, unless otherwise stated. Clinical isolates were collected from patients with wound infection at Tan Tock Seng Hospital, Singapore, under the supervision of Dr Timothy Mark Barkham Sebastian (Laboratory Medicine) and identified as *Enterococcus faecalis* strains by mass spectrometry. For the OG1RF strain transformed with P*c@*-GFP, BHI was supplemented with spectinomycin (120 µg/ml). Growth curves were generated by diluting overnight cultures 1:100 in IMDM (Gibco) and adding 100 µl of the diluted cultures in triplicate to a 96-well plate. The plate was then incubated for 18 h at 37°C in a plate reader (TECAN), with absorbance recorded at 600 nm to quantify growth and at 439 nm to detect acidification every 20 min in ambient air conditions without shaking.

### Human Neutrophil Isolation

Whole blood was collected from healthy adult volunteers after written informed consent under a protocol approved by the Nanyang Technological University Institutional Review Board (IRB-2020-06-005). Eligible participants were 21–65 years old, male or female. Participant-level age, sex, gender identity, and ethnicity were not recorded, and blood samples were transferred to the research laboratory without donor identifiers. Neutrophils from EDTA-treated blood were isolated using MACSxpress Whole Blood Neutrophil Isolation Kit (Miltenyi Biotec, Cat. No. 130-104-434) according to the manufacturer’s instructions. Briefly, following the magnetic separation of neutrophils from other white blood cells and erythrocytes, neutrophils were centrifuged at 500 x g for 5 min at 4°C and the pellet was resuspended in fresh IMDM media and cells were enumerated.

### Mammalian cell culture

The human promyelocytic leukaemia cell line HL-60 (ATCC CCL-240) was maintained in IMDM medium supplemented with 20% heat-inactivated fetal bovine serum (FBS; Gibco) at 37°C in a humidified incubator with 5% CO₂. Cells were cultured in suspension and maintained at a density between 2 × 10⁵ and 1 × 10⁶ cells/mL. For neutrophil-like differentiation, HL-60 cells were seeded at 6 × 10⁵ cells/mL and treated with 70 mM N,N-dimethylformamide (DMF; Sigma-Aldrich). Cells were differentiated (dHL-60) for 6-7 days before use in experiments.

### *In vitro* biofilm infection assay

For biofilm formation, overnight BHI cultures were washed and resuspended in an equal volume of IMDM. Resuspended cultures (500 μL) of 1 × 10^9^ CFU/mL were seeded into 24-well plates and incubated for 30 min at 37°C with 5% CO₂ to allow bacterial attachment. For immunofluorescence experiments, wells contained round glass coverslips pre-coated with 0.0001% poly-L-lysine and 250 μg/mL human fibrinogen (Sigma-Aldrich). After attachment, wells were washed twice with PBS to remove non-adherent bacteria, fresh IMDM was added, and biofilms were grown for 24 h at 37°C with 5% CO₂. IMDM was supplemented with spectinomycin (120 µg/ml) when growing WT P*c@*-GFP.

For biofilm infection experiments, the medium of the 24-h biofilm was replaced with 500 μL fresh IMDM containing neutrophils or dHL-60 cells at the required concentration, together with the indicated compounds, and incubated for 4 h at 37°C with 5% CO₂ unless otherwise specified. The biofilm CFU/ml was 1-5 × 10^8^ CFU/mL at the start of the infection for both WT and Δ*ldh* biofilms. For immunofluorescence experiments, neutrophils were added at 0.5 × 10⁶ cells/well. For biofilm supernatant experiments, 24-h biofilms were washed and incubated in fresh IMDM for an additional 4 h. Supernatants were collected, centrifuged at 8000 rpm for 5 min at RT, and filter-sterilized through a 0.2-μm filter to remove residual bacteria. For fixed biofilm experiments, 250 μL of the 500 μL culture medium per well with 24-h biofilm was gently replaced with 250 μL of 8% paraformaldehyde (PFA), yielding a final concentration of 4% PFA, and incubated for 30 min at 37°C. Fixed biofilm was washed twice in PBS, before adding neutrophils. In parallel, a CFU control was included for each experiment to confirm loss of biofilm viability after fixation, with CFU counts being consistently 0. For planktonic bacteria experiments, overnight cultures in BHI were washed and resuspended in IMDM. Bacteria were added to 500 μL fresh IMDM containing neutrophils (1 × 10⁶ cells/ml) at a final concentration of 1 × 10^9^ cfu/ml (MOI 1000), 1 × 10^8^ cfu/ml (MOI 100), 1 × 10^7^ cfu/ml (MOI 10). Where indicated, we added PMA (100 nM; Sigma Aldrich), ionomycin (4 μM; Sigma Aldrich), lactic acid (20 mM; Sigma Aldrich), sodium L-lactate (1, 5, or 20 mM; Sigma Aldrich), sodium D-lactate (1, 5, or 20 mM; Sigma Aldrich), and cytochalasin D (30 μM; abcam, Cat. No. ab143484), pan-MCT inhibitor 7ACC2 (20 μM, HY-D0713, MedChemExpress), pan-MCT inhibitor α-Cyano-4-hydroxycinnamic acid (CHC; 40 μM, HY-107641, MedChemExpress), MCT-1 inhibitor AZD3965 (25 nM, HY-12750, MedChemExpress), MCT-1 inhibitor MCT1-IN-2 (100 nM, HY-18974, MedChemExpress). Where indicated, neutrophils were also pre-incubated with compounds for 30 min before addition to biofilms. IMDM was adjusted from pH 7.4 to pH 6.4 or 5.4 by titration with 1 M HCl.

### Immunofluorescence staining

To preserve NET structures at the infection endpoint, 250 μL of the 500 μL culture medium per well was gently replaced with 250 μL of 8% PFA, yielding a final concentration of 4% PFA, and samples were fixed for 15 min at 37°C. Fixed neutrophils attached to round glass coverslips were then permeabilized with 0.5% Triton X-100 for 5 min and blocked in PBS containing 5% goat serum, 5% BSA, and 0.05% Triton X-100 for 30 min at RT. Samples were incubated overnight at 4°C with rabbit anti-histone H3 (citrulline R2 + R8 + R17) (1:500; Cat. No. ab5103, abcam, RRID: AB_304752) or goat anti-human/mouse myeloperoxidase (1:40; Cat. No. AF3667, R&D Systems, RRID:AB_2250866) in staining buffer (1% goat serum, 1% BSA, 0.05% Triton X-100 in PBS), washed three times, and incubated with fluorophore-conjugated secondary antibodies goat anti-rabbit IgG Alexa Fluor 488 (1:1000; Cat. No. A11034, Thermo Fisher Scientific, RRID:AB_2576217) or donkey anti-goat IgG Alexa Fluor 568 (1:1000; Cat. No. A11057, Thermo Fisher Scientific, RRID:AB_2534104) respectively for 1 h at RT in the dark. F-actin was stained with Alexa Fluor 568-phalloidin (1:500; Thermo Fisher Scientific, Cat. No. A12380) for 30 min at RT and nuclei with 4’,6-diamidino-2-phenylindole (DAPI) (1:500, stock concentration 5 mg/mL, Thermo Fisher Scientific) for 5 min at RT. Sections were mounted in antifade medium and stored in -20°C until imaging.

### Microscopy and image analysis

Stained neutrophil samples were imaged using a Zeiss Axio Observer Z1 inverted widefield microscope at 20× magnification, unless otherwise specified, using ZEN 2 (blue) software (Zeiss). For each sample and condition, a minimum of 5 nonoverlapping fields of view (FOVs) was acquired. When overall cell density was low, imaging continued, with up to 10 FOVs acquired. For each experimental condition, 5–10 FOVs were analyzed, with two exceptions: four FOVs were analyzed for donor 2 under the PMA condition in Fig. 2B and for donor 1 under the 5 mM D-lactate condition in Fig. 2F. NET formation was quantified by counting decondensed nuclei, H3Cit^+^ nuclei, cells releasing NETs, and total nuclei using the Cell Counter plugin in ImageJ (version 1.54p). Representative neutrophil nuclear morphologies used for NETosis classification are shown in **Supplementary Figure S1**. Representative image panels were cropped from the original images to a final field size of 50 × 50 μm. Quantification was performed on full size images using the following formulas: Decondensed nuclei (%) = (Number of decondensed nuclei / Total nuclei) × 100, H3Cit^+^ nuclei (%) = (Number of H3Cit^+^ nuclei / Total nuclei) × 100, and NETosis (%) = (Number of cells releasing NETs / Total nuclei) × 100. Neutrophils with intracellular bacteria were quantified based on microscopy images using the following formula: Neutrophils with engulfed bacteria (%) = (Number of neutrophils with intracellular bacteria / Total neutrophils) × 100. Z-stacks of neutrophil samples were acquired on a Carl Zeiss LSM 780 laser scanning confocal with a 63× oil immersion lens using ZEN 2.3 SP1 FP3 (black) software (Zeiss). Maximum intensity projections and orthogonal views were generated from Z-stacks using ImageJ (version 1.54p). Image acquisition settings were kept constant across all conditions. Linear adjustments of brightness and contrast were applied exclusively to the DAPI channel to improve visualization of neutrophil nuclei and NETs. No other image modifications were performed. All microscopy images were acquired at the NTU Optical Bio-Imaging Centre (NOBIC) imaging facilities at SCELSE.

### Cell viability assay

Cell viability was assessed by propidium iodide (PI; Thermo Fisher Scientific) staining. Briefly, neutrophils were seeded at 5 × 10^4^ cells per well in a 96-well plate and exposed to indicated experimental conditions for 4 h at 37°C and 5% CO₂. Following treatment, PI was added directly to each well at a final concentration of 5 μM and incubated for 15 min at RT, protected from light. Fluorescence was measured using a plate reader with excitation at 535 nm and emission at 617 nm. Background fluorescence from cell-free wells containing medium and PI was subtracted from all readings. As a positive control for membrane permeabilization, neutrophils in medium were treated with 0.1% Triton X-100 for 5 min at RT before PI addition. Values were normalized to the positive control condition, which corresponded to 0 % viability.

### Biofilm CFU quantification

24-h biofilms grown in 24-well plates as described for the *in vitro* biofilm infection assay were gently washed twice with PBS to remove non-adherent bacteria. Fresh IMDM was added containing human neutrophils (2 × 10⁶ cells/well) where indicated, with control wells receiving IMDM alone. Where indicated, neutrophils were pre-treated for 30 min with cytochalasin D (CytoD; 30 µM), and incubated with biofilm for 4 h at 37°C with 5% CO₂, under the continued presence of cytochalasin D to assess the contribution of phagocytosis to biofilm control. At 0 or 4 h post-exposure, biofilms were detached with scraping, serially diluted in PBS and plated on BHI agar. CFU were enumerated after overnight incubation at 37°C.

### Flow cytometry

To quantify bacterial engulfment, dHL-60 cells (1 × 10⁶) were incubated for 1 h with WT Pcfb-GFP biofilms in IMDM supplemented with spectinomycin (120 μg/mL). Where indicated, sodium L-lactate, sodium D-lactate, or lactic acid was added at 20 mM. Cytochalasin D (30 μM) was used as a negative control for engulfment; cells were pre-incubated with cytochalasin D for 30 min before addition to biofilms, and cytochalasin D was maintained throughout the subsequent 1-h incubation. Following infection, cells were scraped, pelleted (500g, 5 min, 4°C) and resuspended in 94 µL flow cytometry staining buffer (FCSB; 1% BSA and 0.1% Sodium Azide). Anti-human Fc block (1 μL/sample; Cat. No. 564219, BD Pharmingen, RRID: AB_2728082) was added for 20 min at 4°C. Cells were then stained with anti-human CD45-Alexa Fluor 700 (5 μL/sample; Cat. No. 56-9459-42, Thermo Fisher Scientific, RRID: AB_2574511) in a final volume of 100 μL for 30 min at 4°C in the dark. Samples were washed by adding 400 μL FCSB, centrifuged at 500 × g for 5 min at 4°C, and fixed by resuspending the pellet in 100 μL of 4% PFA for 15 min at 4°C in the dark. After fixation, samples were washed once more with 400 μL FCSB, centrifuged at 500 × g for 5 min at 4°C, and resuspended in 200 μL FCSB. Samples were stored at 4°C in the dark until acquisition. A total of 10,000 events were acquired per sample, and bacterial engulfment was quantified as the mean FITC fluorescence intensity of CD45⁺ events, corresponding to GFP-expressing intracellular bacteria. Data were normalized to the control condition containing dHL-60 cells incubated with WT Pcfb-GFP biofilms only, which corresponded to 100% engulfment.

To quantify neutrophils in infected murine wounds, murine wound beds were excised, weighed, and minced in 500 µL of digestion buffer (0.1 mg/mL Liberase in DMEM), followed by incubation at 37 °C for 60 min with gentle rocking. After incubation, DMEM (2 × 500 µL) was added, and samples were filtered through a 70-µm strainer. Filtrates were centrifuged at 500 × g for 5 min at 4 °C, and cell pellets were resuspended in 200 µL of FCSB. For staining, 100 µL per sample were used. Fc receptors were blocked with TruStain FcX PLUS anti-mouse CD16/32 antibody (clone S17011E; BioLegend, Cat. No. 156604, RRID: AB_2783138) for 20 min at 4°C, followed by staining with Ly6G-APC (Cat. No. 127614, BioLegend, RRID: AB_2227348), CD45-BV510 (Cat. No. 103138, BioLegend, RRID: AB_2563061), and CD11b-PE (Cat. No. 101208, BioLegend, RRID: AB_312791) at 1:100 for 30 min at 4°C in the dark. After incubation, cells were washed and fixed in 4 % PFA for 15 min at 4 °C in the dark, followed by washing and resuspension in 100 µL FCSB. To obtain absolute cell counts, 100 µL of AccuCheck Counting Beads (Thermo Fisher Scientific, Cat. No. PCB100) were added to each sample by reverse pipetting. Compensation was applied using the AbC Anti-Mouse Bead Kit (Thermo Fisher Scientific, Cat. No. A10344), and fluorescence-minus-one (FMO) controls were used to set gating thresholds. A total of 10000 events were collected per sample. Flow cytometry data were analyzed using FlowJo (version 10.7.1). Absolute cell counts were calculated according to the manufacturer’s instructions for AccuCheck Counting Beads. All samples were acquired using a BD LSRFortessa X-20 flow cytometer equipped with 5 lasers (488 nm, 535 nm, 633 nm, 405 nm, 355 nm).

### Isothermal microcalorimetry

dHL-60 cells or human neutrophils were resuspended in fresh IMDM or biofilm supernatant at 10 × 10^6^ cells/ml and 200 µL (2 × 10^6^ cells) was loaded per condition, in duplicate, into calVial Inserts (SymCel, Sweden). The calVial Inserts were placed into a calPlate containing titanium vials (SymCel, Sweden) and incubated in a SymCel calScreener (SymCel, Sweden) for 4 h at 37°C to measure cellular heat production. After incubation, the first 1.5 h of recording, corresponding to the thermal equilibration phase during which the signal is not reliable for analysis, was excluded from the dataset. A blank sample containing only IMDM was included as negative control.

### Targeted metabolomics

Metabolite extraction and targeted metabolomics analyses of metabolites in central carbon metabolism followed the published reports with modifications (*71*, *72*). Briefly, cell cultures were harvested and rapidly quenched, and metabolites were extracted using acetonitrile:methanol:water (2:2:1) through three freeze-thaw cycles. After centrifugation, the supernatant was collected and evaporated to dryness in a vacuum evaporator, and the dry extracts were redissolved in 100 µL of 98:2 water/methanol for liquid chromatography-mass spectrometry (LC-MS) analysis. The targeted LC-MS/MS analysis was performed with Agilent 1290 ultrahigh pressure liquid chromatography system coupled to a 6495 Triple Quadrupole mass spectrometer (Agilent Technologies). Chromatographic separation of metabolites in central carbon metabolism was achieved by using Phenomenex RezexTM ROA-Organic Acid H+ (8%) column (2.1×100 mm, 3 µm) and the compounds were eluted at 40°C with an isocratic flow rate of 0.3 mL min-1 of 0.1% formic acid in water. Compounds were quantified in multiple reaction monitoring (MRM) mode. Electrospray ionization was performed in negative ion mode with the following source parameters: drying gas temperature 250°C with a flow of 10 L min-1, nebulizer gas pressure 40 psi, sheath gas temperature 350°C with a flow of 11 L min-1, nozzle voltage 500 V, and capillary voltage 2,500 V. Data acquisition and processing were performed using MassHunter software (Agilent Technologies).

### ATP quantification

dHL-60 cells or human neutrophils were resuspended in fresh IMDM or the indicated biofilm supernatants and reagents, and seeded into black 96-well plates at 5 × 10^4^ cells/well in a total volume of 100 µL. Cells were incubated at 37°C in 5% CO₂ for 4 h. ATP levels were then quantified using the CellTiter-Glo 2.0 Cell Viability Assay (Promega) according to the manufacturer’s instructions. Briefly, an equal volume of CellTiter-Glo 2.0 reagent was added directly to each well, contents were mixed with shaking for 2 min to induce cell lysis, followed by incubation at RT for 10 min. Luminescence, proportional to ATP content, was measured using a plate reader (TECAN). Background-subtracted luminescence values were normalized to the media control and expressed as log2 fold change in the plots.

### Intracellular pH quantification

dHL-60 cells (5 × 10^5^/condition) were loaded with the pH-sensitive fluorescent dye BCECF-AM (10 μM; Thermofisher) in IMDM at 37°C, 5% CO_2_ for 30 min, washed to remove excess dye, and resuspended in fresh IMDM or the indicated biofilm supernatants. After 4 h incubation at 37°C, 5% CO_2,_ fluorescence was measured using a plate reader with dual excitation at 440 nm and 488 nm and emission at 535 nm. To generate a standard curve, BCECF-AM-loaded dHL-60 cells with CCCP (2 μM) for 30 min in IMDM, following addition of IMDM adjusted to defined pH values. Fluorescence was then measured using the same settings (440/488 nm excitation, 535 nm emission). The 488/440 nm fluorescence ratio was plotted against the corresponding pH values to generate a standard curve, which was used to interpolate intracellular pH in experimental samples.

### NADP+/NADPH quantification

Intracellular total NADP+/NADPH levels in neutrophils were quantified using the NADP/NADPH Assay Kit (ab65349, abcam) according to the manufacturer’s instructions. Briefly, 2 x 10^6^ neutrophils/well were incubated for 4 h with IMDM, WT or Δ*ldh* biofilm supernatant in a 24-well plate. Neutrophils were collected with scraping, washed with cold PBS, and lysed with lysing matrix B (MP Biomedicals, Cat. No. 116923050) in 600 µL extraction buffer II by beating for 40 s at 6 m/s. Lysates were clarified by centrifugation. 100 µL of each sample was added to a clear 96-well plate, followed by 100 µL Master Reaction Mix and 10 µL Developer Solution II. For each sample, background OD_450nm_ values from wells containing sample and Background Reaction Mix were subtracted from OD_450nm_ values obtained with the full reaction mix. Background-corrected OD₄₅₀ values were analyzed and presented without normalization.

### Murine wound infection model

Six-week-old male C57BL/6J mice were used in accordance with NTU Institutional Animal Care and Use Committee protocols A19061 and A24066. Sample sizes were based on previous experiments using this murine wound infection model (*11*). Mice were randomly allocated to the WT- or Δ*ldh*-infected groups. Investigators were not blinded to group allocation during the experimental procedures or outcome analyses. Animals were excluded if the wound dressing became detached or was lost before the scheduled endpoint. In addition, mice were excluded if no bacteria were recovered from the wound, indicating that infection had not been established. Mice were anesthetized with 2% isoflurane, and dorsal hair was removed. The skin was disinfected 2-3 times with 70% ethanol before generating a 6-8 mm full-thickness excisional wound using a 6 mm biopsy punch. Wounds were inoculated with 10 µL bacterial suspension (∼1-2 × 10⁶ CFU) in PBS and 3M Tegaderm Transparent Film Dressing (Cat. No. 1622W) was applied for 1 or 7 days. For 7-day infections, Tegaderm was replaced every 2 days or as needed. Buprenorphine was injected (0.05–0.2 mg/kg s.c.) for analgesia as required. At the experimental endpoint, mice were euthanized with CO_2_ inhalation and the entire wound bed was excised and kept on ice before processing for protein isolation or flow cytometry analysis.

### Western Blot

Infected wound beds were transferred to 2 mL Lysing Matrix M tubes containing 1 mL PBS and homogenized for 5 rounds at 4m/s for 20s, keeping the sample on ice after each round. Samples were centrifuged, and the supernatant was collected for protein analysis. Protein concentration was determined using the BCA Protein Assay Kit (Thermo Fisher Scientific) according to the manufacturer’s instructions. 16 µg of protein per sample was separated on 4-12% NuPAGE Bis-Tris gels at 120 V for ∼45 min and transferred to PVDF membranes using the iBlot2 Gel Transfer Device (Thermo Fisher Scientific). Post-transfer Coomassie blue staining was applied on the gels to evaluate protein loading across lanes. Membranes were blocked in 5% BSA in TBS-T (0.1% Tween-20) for 1 h at RT and incubated with rabbit anti-histone H3 (citrulline R2 + R8 + R17) (1:1000; Cat. Nr. ab5103, abcam, RRID: AB_304752) or rabbit anti-histone H3 (1:2000; Cat. Nr. ab1791, abcam, RRID: AB_302613) diluted in 2% BSA/TBS-T overnight at 4°C. After washing (3 × 5 min in TBS-T), membranes were incubated with goat anti-Rabbit IgG (H+L) HRP-conjugated antibody (1:5000, Cat. Nr. 31460, Invitrogen, RRID: AB_228341) for 1 h at RT, followed by additional washes (3 × 5 min in TBS-T). Protein bands were visualized using SuperSignal West Femto Maximum Sensitivity Substrate (Thermo Fisher Scientific) and imaged in Amersham ImageQuant 800. Loading controls were run on the same blot. For reprobing, membranes were stripped with Restore PLUS Western Blot Stripping Buffer (30 min, 50°C). Band intensities were quantified using ImageJ (version 1.54p). Coomassie blue stain and full membrane images are included in the Source Data File.

### ELISA

Wound lysates from 1 and 7 dpi were stored at -80 °C until assessment by the mouse CCL2/JE/MCP-1 (Cat. No. # DY479, R&D Systems) and the mouse CXCL1/KC quantikine ELISA Kits (Cat. No. # DY453, R&D Systems) according to manufacturer’s recommendations. All samples were assessed using the same kit lot and at the same time to avoid inter-assay variability.

### Statistical Analysis

Unless otherwise stated, N denotes the number of independent biological experiments. For primary human neutrophil assays, each biological replicate comprises an independently performed experiment using a separate blood collection and neutrophil isolation. For dHL-60 and bacterial assays, biological replicates were independently performed experiments using separately prepared cultures. Technical replicate wells and microscopy fields were combined within each experiment to generate one value per biological replicate and were not treated as independent observations. For animal studies, N denotes the number of independent animal experiments and n denotes the number of individual mice. Individual mice are defined as biological replicates in statistical analysis. Exact replicate numbers are provided in the respective figure legends. Prism version 11.0.2 (GraphPad, USA) was used for all statistical analyses. Unless otherwise stated, comparisons between two independent groups were performed using two-tailed unpaired t-tests or two-tailed Welch’s t-tests, as specified in the figure legends. Several multi-condition neutrophil datasets were heteroskedastic and contained zero-variance groups, precluding the use of one-way ANOVA or Welch’s ANOVA. For these datasets, statistical analysis was restricted to the hypothesis-driven pairwise comparisons shown in the figures, using two-tailed paired t-tests. For LC-MS analysis, technical replicate peak areas were averaged within each biological replicate before statistical analysis. Metabolites that were not detected in any biological replicate within a condition were classified as undetected. Statistical comparisons were performed on biological-replicate mean peak areas using separate two-tailed Welch’s t-tests for WT versus media and Δldh versus media. Comparisons were not performed when one or more biological replicates in either group had an undetected value and were reported as not applicable (NA). Statistical significance was defined as P < 0.05.

## ACKNOWLEDGEMENTS

We thank Tim Barkham and Kelvin Chong for providing and sequencing clinical *E. faecalis* isolates TTSHW-EF35, TTSHW-EF37. We also thank SCELSE members Kenneth Ng Kok Fei and Audrey Tan Mei Hui for technical assistance in experiment preparation, Sharan Jagdish Prakash for providing training on the calScreener, Claudia J. Stocks for the support in coordinating volunteer recruitment and managing ethics-related logistics for this study, Jessica Alison Taylor, Sujatha Srinivas, and Karen Butina for insightful discussions in statistical analysis. We also thank Alain Filloux (SCELSE) for the institutional support that enabled the completion of this work.

## AUTHORS CONTRIBUTIONS

Conceptualization: H.A., K.A.K.

Methodology: H.A., R.A.G.D.S., C.L., S.L.W., K.A.K.

Formal Analysis: H.A., R.A.G.D.S., G.K.X.W., C.L., S.L.W., K.A.K.

Investigation: H.A., R.A.G.D.S., G.K.X.W., C.L., L.M.N., R.J.W.T., C.J.Y.N.

Resources: H.A., S.L.W., K.A.K.

Writing - Original draft preparation: H.A.

Writing-Review and editing: H.A., R.A.G.D.S., G.K.X.W., C.L., L.M.N., R.J.W.T., C.J.Y.N., S.L.W., K.A.K.

Visualization: H.A.

Supervision: H.A., S.L.W., K.A.K.

Project administration: H.A., S.L.W., K.A.K.

Funding Acquisition: H.A., S.L.W., K.A.K.

## DATA, MATERIALS, AND CODE AVAILABILITY

All data needed to evaluate the conclusions in the paper will be present in the Supplementary Materials and relevant repositories upon publication.

## TABLE OF CONTENTS FOR SUPPLEMENTARY MATERIALS

**Fig. S1.** Representative neutrophil nuclear morphologies used for NETosis classification

**Fig. S2.** Ionomycin-mediated NETosis in human neutrophils is not completely inhibited by *E. faecalis* biofilm

**Fig. S3**. *E. faecalis* inhibits NETosis via lactic acid-mediated extracellular acidification

**Fig. S4.** Effect of *E. faecalis* clinical isolates on NETosis

**Fig. S5.** L- and D-lactate do not induce NETosis in human neutrophils

**Fig. S6.** *E. faecalis*-derived lactic acid suppresses glycolysis, the TCA cycle, and the oxidative pentose phosphate pathway in dHL-60 cells

**Fig. S7.** MCT inhibitors partially inhibit PMA-induced NETosis

**Fig. S8**. Δ*ldh* biofilm inhibits PMA-induced NETosis in neutrophils extracellularly

**Fig. S9.** Addition of L- or D-lactate to Δ*ldh* biofilms does not induce NETosis in human neutrophils

**Fig. S10.** PFA-fixed Δ*ldh* biofilm suppresses NETosis

**Table S1.** Bacterial strains

**Supplementary Data File 1.** LC-MS analysis of primary human neutrophils exposed to media, WT, or Δ*ldh* biofilm supernatant

**Supplementary Data File 2.** LC-MS analysis of dHL-60 cells exposed to media, WT, or Δ*ldh* biofilm supernatant

