## Supplementary Materials for "*Enterococcus faecalis* biofilm rewires neutrophil metabolism to suppress antimicrobial activity"

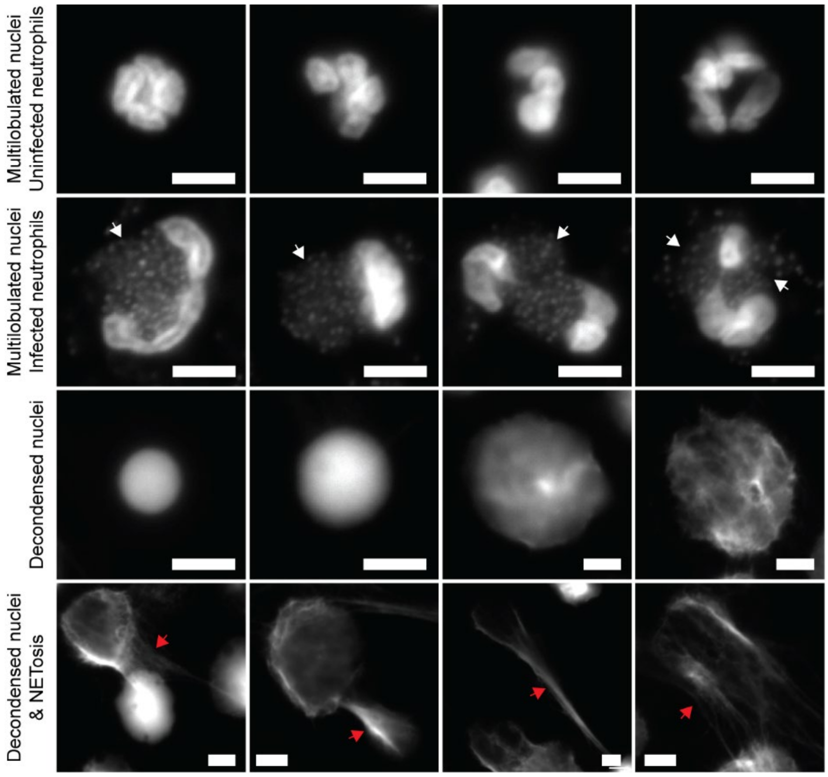


**Supplementary Figure S1. Representative neutrophil nuclear morphologies used for NETosis classification.** DAPI-stained fluorescence microscopy images showing the morphological criteria used to classify neutrophil nuclei. Images were captured using a Zeiss Axio Observer Z1 inverted widefield microscope with a 63× oil immersion lens. Uninfected neutrophils with compact or loose multilobulated nuclei were classified as non-decondensed nuclei. In infected neutrophils, multilobulated nuclei may appear morphologically altered because intracellular bacteria (white arrows) occupy cellular volume and can displace or separate nuclear lobes across different focal planes. Although these nuclei may deviate from the appearance of uninfected neutrophil nuclei, they were still classified as non-decondensed nuclei when nucleus lobulation was retained. Neutrophils showing loss of multilobulated nuclear structure, rounding, and nuclear swelling were classified as having decondensed nuclei. Cells with decondensed nuclei associated with extracellular DNA fibers were classified as nuclei releasing NETs. Red arrows = NETs, white arrows = bacteria, scale bar = 5 µm.

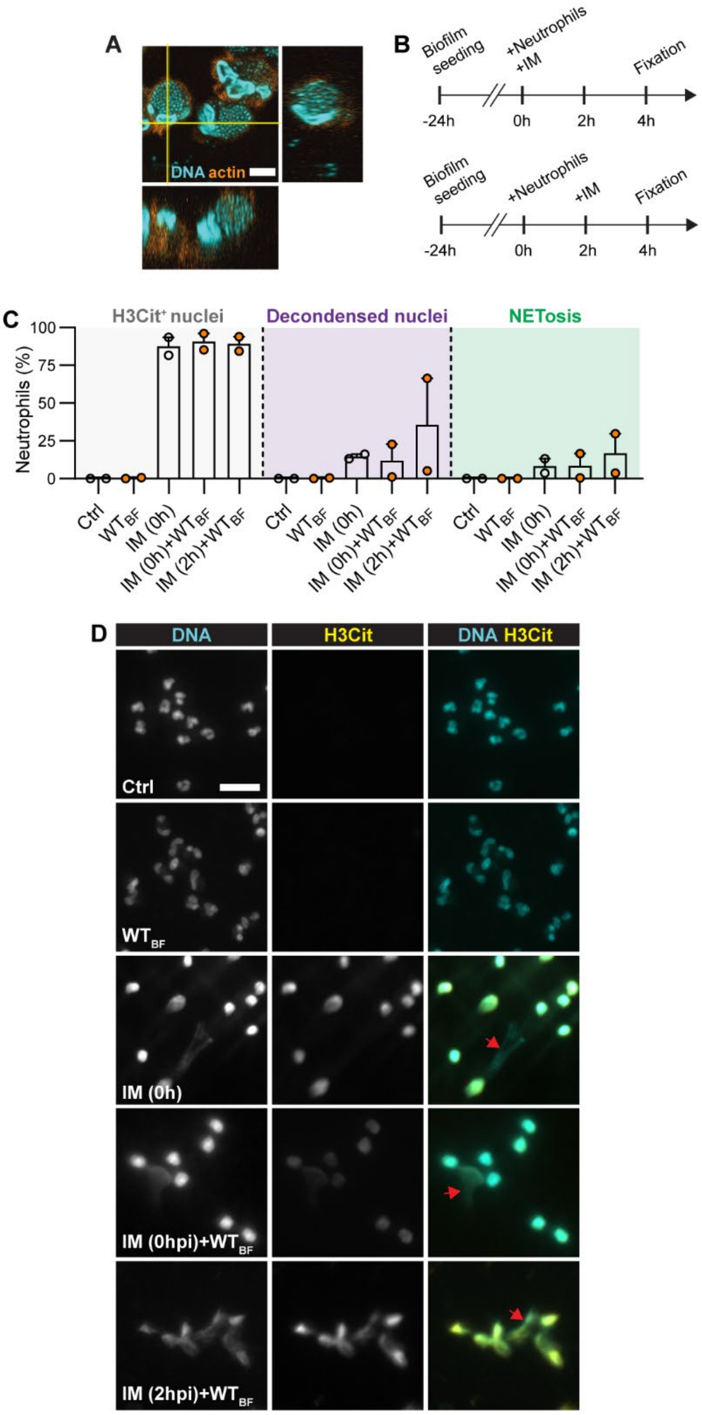


**Supplementary Figure S2. Ionomycin-mediated NETosis in human neutrophils is not completely inhibited by *E. faecalis* biofilm. A.** Human neutrophils with internalized *E. faecalis* after exposure to biofilm for 4 h. Representative single section with orthogonal views from a Z-stack, stained for DNA and F-actin. N = 3, Scale = 5 µm. **B.** Experimental timeline for C-D. Biofilm was grown for 24 h, followed by either simultaneous addition of human neutrophils and ionomycin (IM; 4 µM) at 0 h or addition of neutrophils at 0 h and IM at 2 h, followed by fixation at 4 h. **C-D.** Quantification (C) and representative immunofluorescence microscopy images (D) of NETosis hallmarks in neutrophils stained with DAPI and anti-H3Cit antibody. The percentage of H3Cit⁺ nuclei, decondensed nuclei, and NET-releasing cells (NETosis) was quantified from N = 2 independent experiments and is shown as mean ± SEM. Scale bar = 20 µm; red arrow = NETs. Ctrl = uninfected & unstimulated neutrophils in media. The control sample from one donor shown in panel C is the same control samples in Fig. S10A, as both panels were generated from neutrophils deriving from the same blood donation.

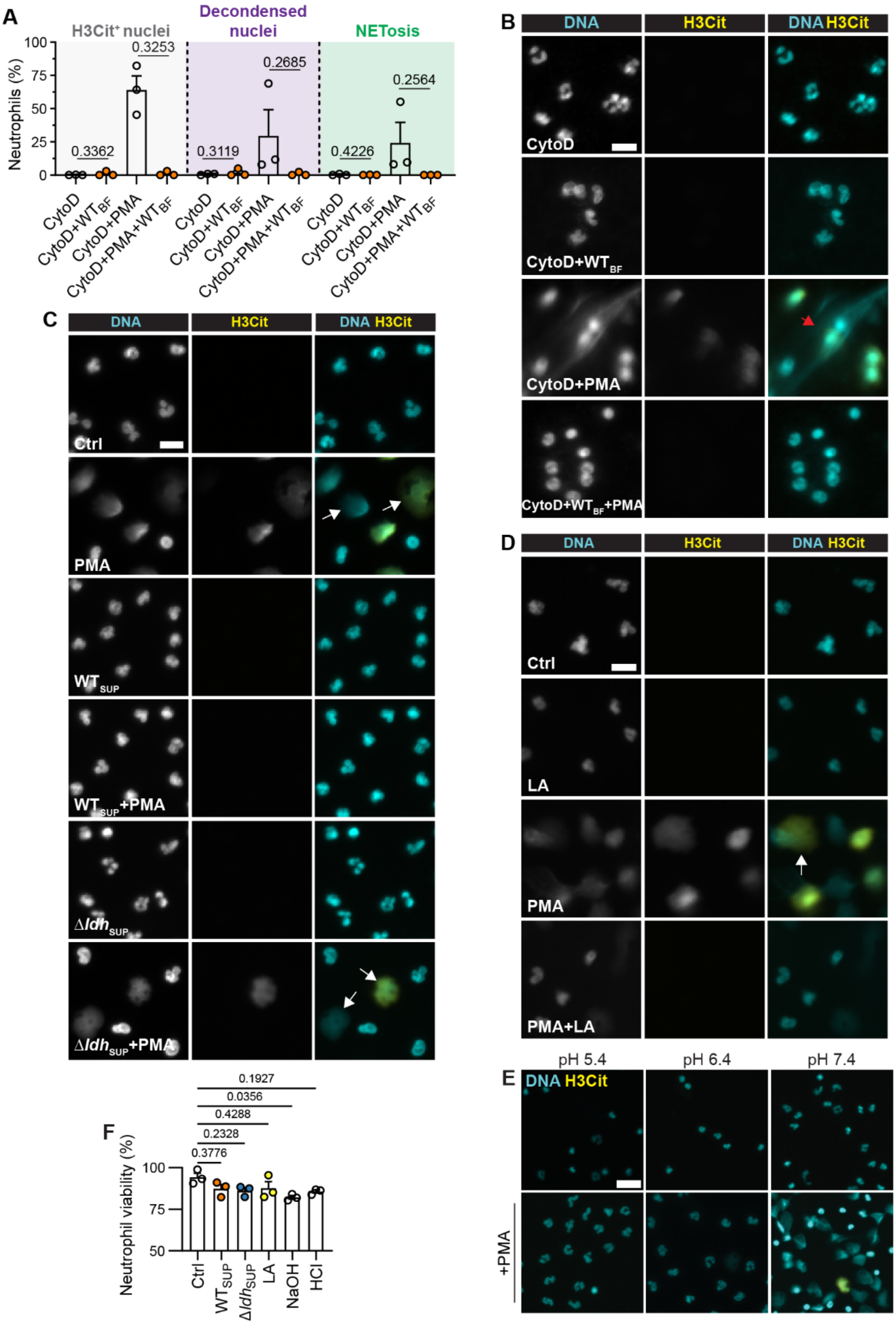


**Supplementary Figure S3.** ***E. faecalis*** **inhibits NETosis via lactic acid-mediated extracellular acidification. A.** Quantification of NETosis hallmarks from immunofluorescence microscopy images. Mean (%) citrullinated histone H3 stained nuclei (H3Cit$^+$), decondensed nuclei, and NETs release (NETosis) from N = 3 independent experiments are shown, error = SEM. Human neutrophils were pre-incubated with Cytochalasin D (30 µM; CytoD) for 30 min prior to addition to the biofilm, stimulated with PMA at 0 h and incubated with *E. faecalis* OG1RF biofilm for 4 h in the continued presence of CytoD. CytoD and cytoD + PMA control samples shown for one donor in panel A are the same control samples shown in Fig. 5A, as one biological replicate for both panels was generated from neutrophils from the same blood donation. Statistical significance was assessed with two-tailed paired t-test. **B-E**. Representative immunofluorescence microscopy images of neutrophils stained with DAPI and anti-H3Cit antibody. N = 3. Fig. S3B corresponds to quantification shown in Fig. S3A, Fig. S3C to Fig. 2A, Fig. S3D to Fig. 2B, Fig. S3E to Fig. 2D. Scale = 10 µm in Fig. S3B-D and 20 µm in Fig. S3E, red arrow = NETs, white arrow = decondensed nuclei. Only merged images shown in Fig. S3E. **F.** Quantification of neutrophil viability by propidium iodide 4 h after exposure to the indicated conditions. Fluorescence values were normalized to a positive control of neutrophils treated with 0.1% Triton X-100. N = 3, error = SEM. Statistical significance was assessed with one-way ANOVA followed by Tukey's multiple comparisons test; $\alpha = 0.05$ with significant values shown in blue.

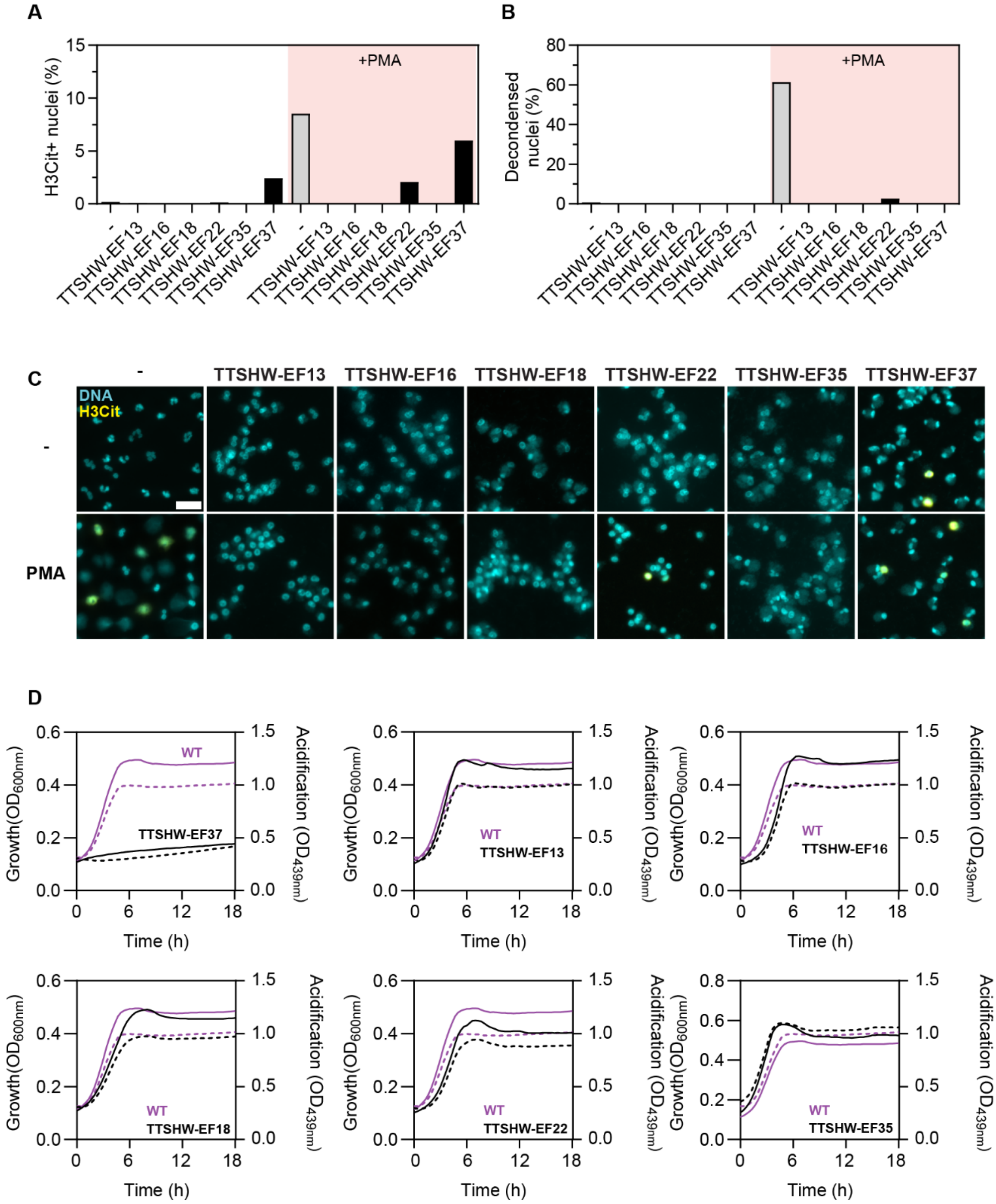


**Supplementary Figure S4. Effect of *E. faecalis* clinical isolates on NETosis. A-C.** Human neutrophils were incubated with *E. faecalis* clinical isolates biofilm for 4 h and stimulated with PMA (100 nM) at 0 h. NETosis was assessed by immunofluorescence microscopy based on nuclear decondensation and citrullinated histone H3 (H3Cit) staining. Mean (%) of H3Cit positive and decondensed nuclei is shown together with representative images from N = 1 experiment. **D.** Growth curves (solid line) based on $OD_{600nm}$ measurements (left Y-axis) and acidification curves (dashed line) based on $OD_{439nm}$ measurements (right Y-axis) of *E. faecalis* clinical isolates in IMDM**.** N = 1.

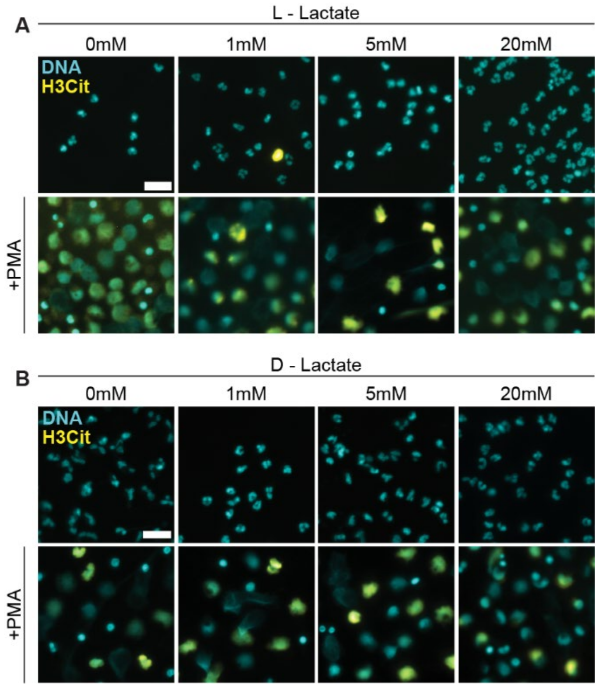


**Supplementary Figure S5. L- and D-lactate do not induce NETosis in human neutrophils. A-B.** Representative immunofluorescence microscopy images of neutrophils stained with DAPI and anti-H3Cit antibody from N = 3, with corresponding NETosis quantification shown in **Fig. 2E-F.** Human neutrophils were incubated for 4 h and stimulated with sodium L- or D-lactate and PMA at 0 h. Scale = 20 μm. Only merged images shown.

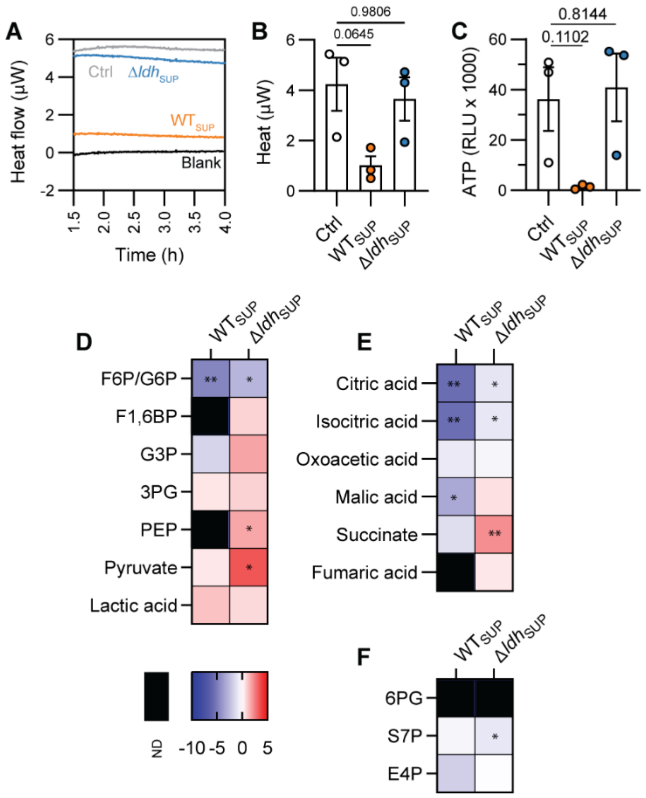


**Supplementary Figure S6.** ***E. faecalis*-derived lactic acid suppresses glycolysis, TCA cycle, and oxPPP in dHL-60 cells. A.** Representative thermogram of dHL-60 cells treated with fresh media (Ctrl), or biofilm supernatant from WT ($WT_{SUP}$) or Δ*ldh* (Δ*ldh*$_{SUP}$) for 4 h. Media alone (blank) was included. Heat measurements start after 1.5 h due to the calScreener initial calibration phase. N = 3. **B.** Mean heat produced by dHL-60 cells at 4 h post-incubation. N = 3, error = SEM. Statistical significance was assessed with paired t-test. **C.** ATP levels in dHL-60 cells after incubation with media alone (Ctrl), $WT_{SUP}$ or Δ*ldh*$_{SUP}$ for 4 h. Data is presented as relative luminescence units, N = 3, error = SEM. Statistical significance was assessed with two-tailed Welch's test. **D-F.** LC-MS heatmaps showing log2 fold changes in metabolite abundance in dHL-60 cells after exposure to $WT_{SUP}$ or Δ*ldh*$_{SUP}$ for 4 h relative to untreated control cells. Metabolites from glycolysis (D), TCA cycle (E), and PPP (F) are shown. N = 3, color scale indicates relative metabolite abundance, with red denoting increased and blue decreased levels relative to control, and black denoting not detected (ND) metabolites. Statistics were calculated on the biological replicate mean peak areas using a two-tailed Welch's t-test to compare $WT_{SUP}$ with media and Δ*ldh*$_{SUP}$ with media for each metabolite. * = P < 0.05, ** = P < 0.01, *** = P < 0.001, **** = P < 0.0001; α = 0.05 with significant values shown in blue.

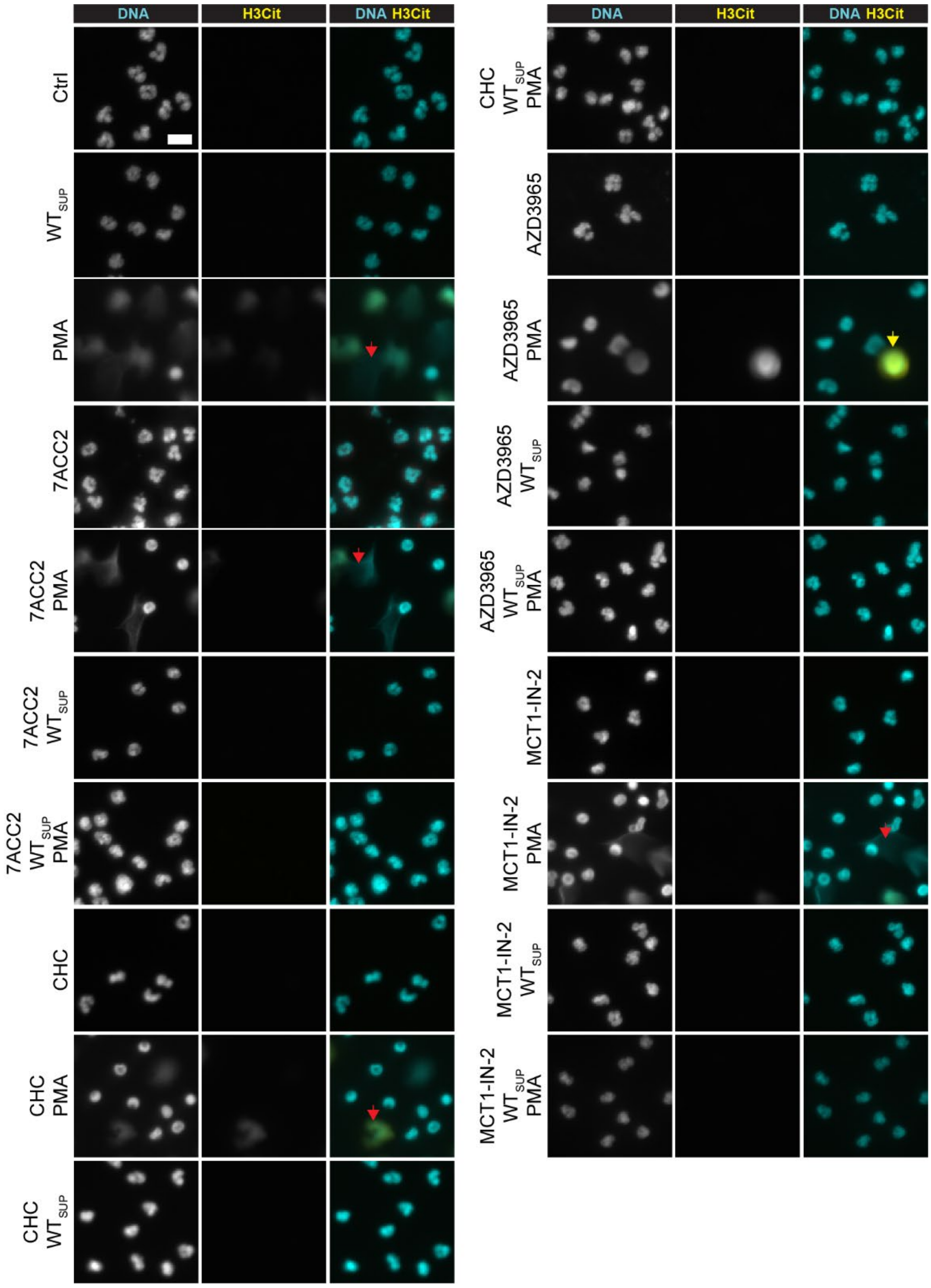


**Supplementary Figure S7. MCT inhibitors partially inhibit PMA-induced NETosis.** Human neutrophils were stimulated with PMA (100 nM) at 0 h and incubated with *E. faecalis* OG1RF WT biofilm supernatant ($WT_{SUP}$) and MCT inhibitors 7ACC2 (20 µM), CHC (40 µM), AZD3965 (25 nM), MCT1-IN-2 (100 nM) for 4 h. NETosis was assessed by immunofluorescence microscopy based on nuclear decondensation and citrullinated histone H3 (H3Cit) staining. Red arrows = NETs, yellow arrow = decondensed nuclei. Representative images are shown from N = 1. Scale = 10 µm.

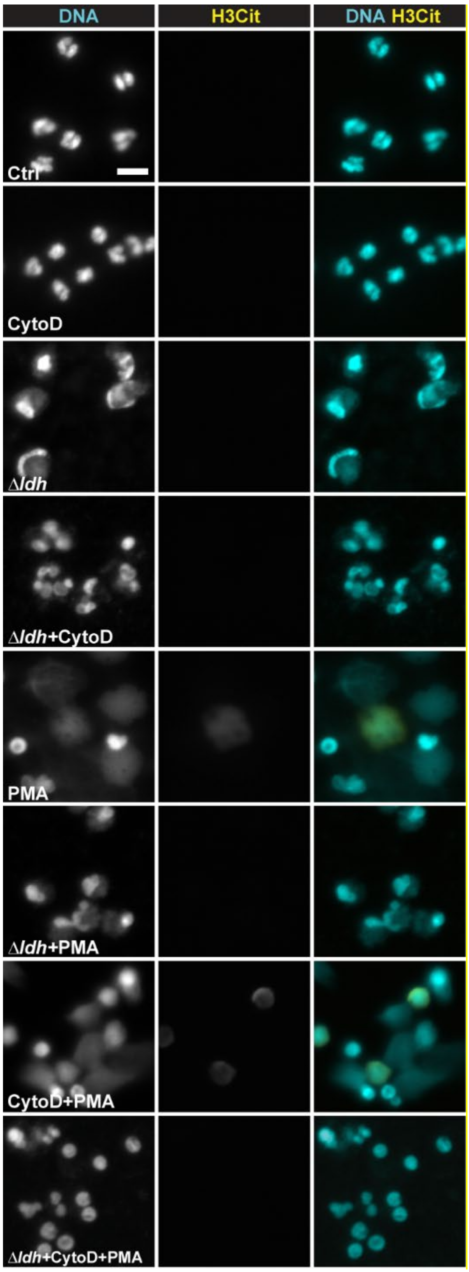


**Supplementary Figure S8. Δ*ldh* biofilm inhibits PMA-induced NETosis in neutrophils extracellularly.** Representative fluorescence microscopy images of human neutrophils incubated with Δ*ldh* biofilm and supplemented with cytochalasin D (cytoD; 30 μM) and PMA (100 nM) as indicated at 4 h post-incubation, corresponding to quantified data in Fig. 5A. Neutrophil nuclei were stained with DAPI and an antibody against citrullinated histone H3 (H3Cit). N = 3, scale = 10 μm.

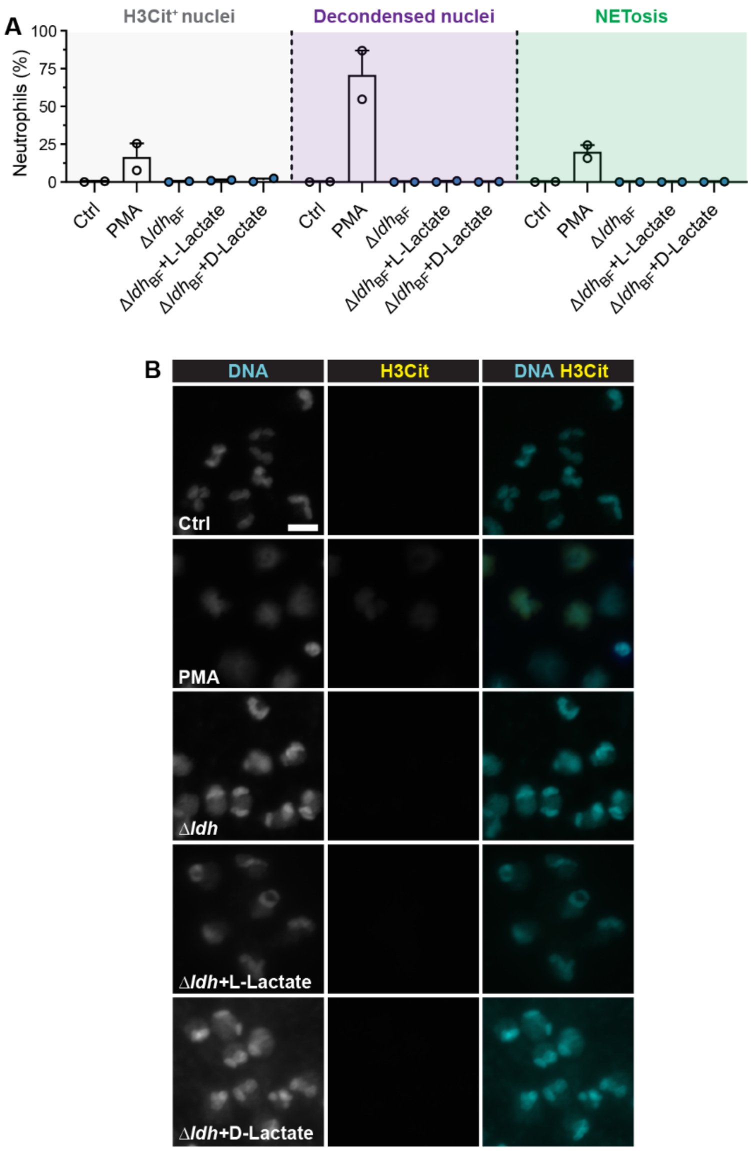


**Supplementary Fig. S9. Addition of L- or D-lactate to Δ*ldh* biofilms does not induce NETosis in human neutrophils. A-B.** Quantification of NETosis hallmarks from immunofluorescence microscopy images. Mean (%) citrullinated histone H3 stained nuclei (H3Cit$^+$), decondensed nuclei, and NETs release (NETosis) from N = 2 independent experiments are shown, alongside representative images. Human neutrophils were supplemented with 20mM of L- or D-lactate at 0 h and incubated with *E. faecalis* OG1RF Δ*ldh* biofilm for 4 h. Ctrl = neutrophils in media, error = SEM, scale = 10 μm.

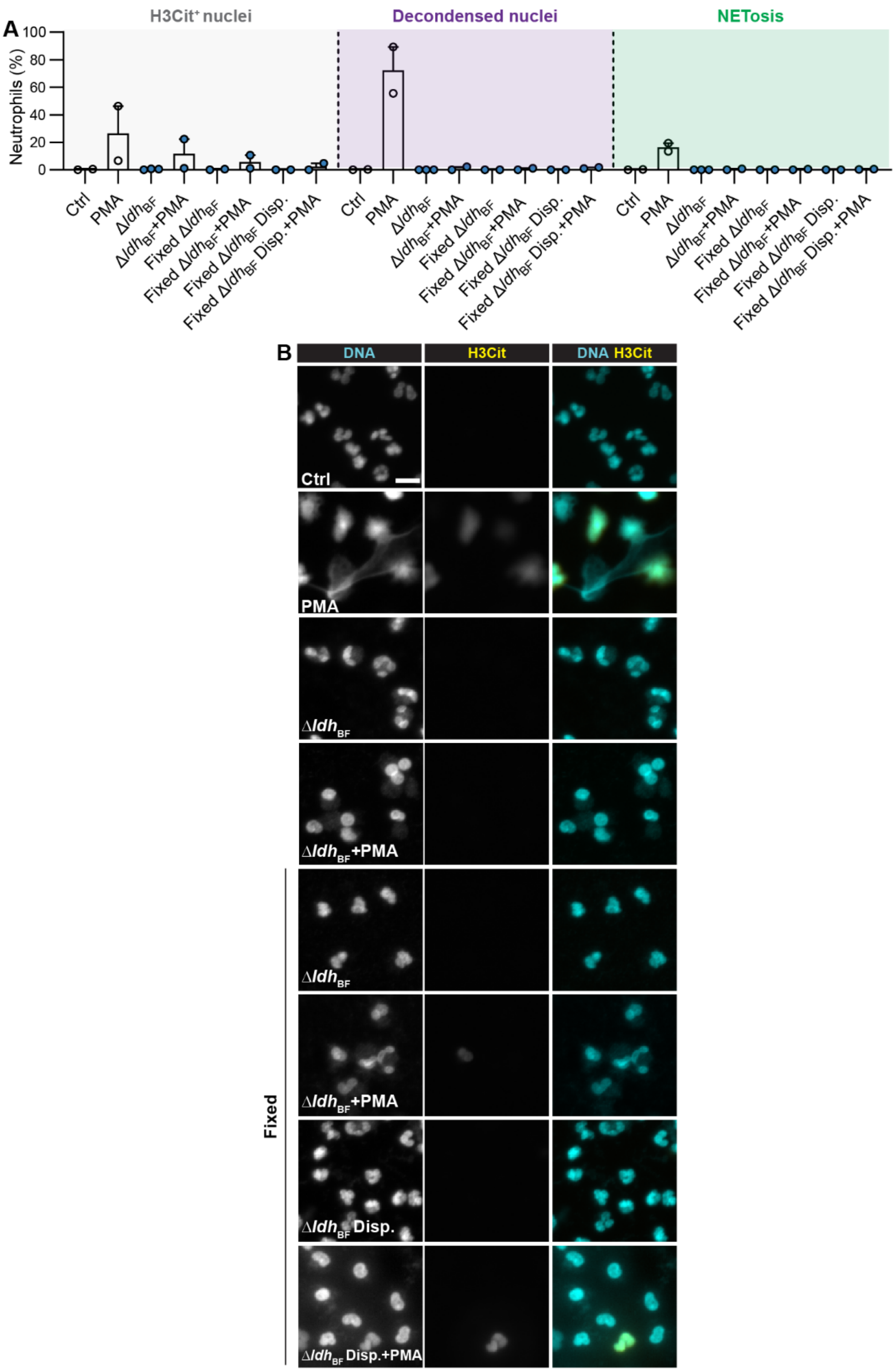


**Supplementary Figure S10. PFA-fixed Δ*ldh* biofilm suppresses NETosis. A.** Quantification of NETosis hallmarks from immunofluorescence microscopy images. Mean (%) citrullinated histone H3 stained nuclei (H3Cit$^{+}$), decondensed nuclei, and NETs release (NETosis) from N = 2 independent experiments are shown, error = SEM. Human neutrophils were incubated with live, PFA-fixed or dispersed (Disp.) Δ*ldh* biofilm (Δ*ldh*$_{BF}$) in the presence of PMA (100 nM) as indicated for 4 h. The control sample shown for one donor in panel A is shared with Fig. S2C, as one replicate from both panels was generated from neutrophils deriving from the same blood donation. **B.** Representative images of neutrophils for S10A with nuclei stained for DAPI and citrullinated histone H3 (H3Cit). N = 2-3, scale = 10 µm.

**Table S1. Bacterial strains**

| Strain | Description | Reference |
|---|---|---|
| OG1RF | *Enterococcus faecalis, $Rif^R$, $Fus^R$* | (1) |
| Δ*ldh* | OG1RF *ldh1* and *ldh2* deletion | (2) |
| WT Pcfb-GFP | OG1RF WT with constitutively expressed Dasher GFP | (3) |
| TTSHW-EF13 | *Enterococcus faecalis* isolate from buttock wound, *cps*+, *fsr*+ | (4) |
| TTSHW-EF16 | *Enterococcus faecalis* isolate from wound swab, *esp*+ | (4) |
| TTSHW-EF18 | *Enterococcus faecalis* isolate from sacral swab, *cyl*+, *cps*+, *fsr*+ | (4) |
| TTSHW-EF22 | *Enterococcus faecalis* isolate from wound swab, *cyl*+, *cps*+, *esp*+ | (5) |
| TTSHW-EF35 | *Enterococcus faecali*s isolate from leg wound, *fsr*+ | This study |
| TTSHW-EF37 | *Enterococcus faecalis* isolate from peripancreatic fluid, *cyl*+, *esp*+ | This study |

[1]Dunny, G. M., Brown, B. L. & Clewell, D. B. (1978) Induced cell aggregation and mating in Streptococcus faecalis: evidence for a bacterial sex pheromone. Proc. Natl. Acad. Sci. U. S. A. 75, 3479–3483.
[2]Silva, R. A. G. da et al. (2026).Enterococcus faecalis-derived lactic acid suppresses macrophage activation to facilitate persistent and polymicrobial wound infections. Cell Host Microbe 34, 245-262.e8
[3]Hallinen, K.M., Guardiola-Flores, K.A., and Wood, K.B. (2020). Fluorescent reporter plasmids for single-cell and bulk-level composition assays in E. faecalis. PLoS One 15, e0232539.
[4]Ch'ng, J. H. et al. (2022). Heme cross-feeding can augment Staphylococcus aureus and Enterococcus faecalis dual species biofilms. ISME Journal 16, 2015–2026.
[5]Stocks CJ, et al. (2026). *Enterococcus faecalis* persists and replicates intracellularly within neutrophils. Infect Immun 94:e00364-25.
